# EHD2-mediated restriction of caveolar dynamics regulates cellular lipid uptake

**DOI:** 10.1101/511709

**Authors:** Claudia Matthäus, Ines Lahmann, Séverine Kunz, Wenke Jonas, Arthur Alves Melo, Martin Lehmann, Elin Larsson, Richard Lundmark, Matthias Kern, Matthias Blüher, Dominik N. Müller, Volker Haucke, Annette Schürmann, Carmen Birchmeier, Oliver Daumke

## Abstract

Epsl5-homology domain containing protein 2 (EHD2) is a dynamin-related ATPase located at the neck of caveolae, but its physiological function has remained unclear. Here, we found that global genetic ablation of EHD2 in mice led to increased fat accumulation. This organismic phenotype was paralleled at the cellular level by increased lipid uptake via a caveolae-, dynamin- and CD36-dependent pathway, an elevated number of detached caveolae and higher caveolar mobility. Furthermore, EHD2 expression itself was down-regulated in the visceral fat of two obese mouse models and obese patients. Our data suggest that EHD2 controls a cell-autonomous, caveolae-dependent lipid uptake pathway and suggest that low EHD2 expression levels are linked to obesity.

## Introduction

Caveolae are small membrane invaginations of the plasma membrane that are abundantly found in adipocytes, endothelial and muscle cells (1). They have been implicated in the regulation of membrane tension (2, 3), in mediating lipid metabolism (4), or in acting as distinct sites for specific and highly regulated signaling cascades such as the endothelial nitric oxide synthase (eNOS)-nitric oxide (NO) pathway (5). The characteristically shaped caveolar bulb has a typical diameter of 50 - 100 nm and is connected to the cell surface via a narrow neck region. The integral membrane protein Caveolin (with three isoforms in human, Cav1-3) and the peripheral membrane protein Cavin (with four isoforms in human, Cavin1-4) build a mesh-like coat around the caveolar bulb (6–10). In addition, BAR domain containing proteins of the PACSIN/syndapin family (PACSIN1-3 in human) participate in the biogenesis of caveolae (11–13).

Loss of Cav1/Cav3 or Cavin1 results in a complete lack of caveolae from the plasma membrane (4, 14, 15). Cav1 KO mice suffer from cardiomyopathy, pulmonary hypertension, endothelium-dependent relaxation problems and defective lipid metabolism (1). In agreement with the latter, Cav1 KO mice are resistant to high fat diet-induced obesity (16) and display smaller white adipocytes and fat pads (17). Furthermore, increased levels of triglycerides and fatty acids are found in blood plasma samples obtained from Cav1 KO mice suggesting a reduced cellular uptake of fatty acids (16). A similar metabolic phenotype was found in mice lacking Cavin1 (4, 18). Conversely, overexpression of Cav1 in adipocytes results in an increased number of caveolae, enhanced fat accumulation, enlarged adipocytes and lipid droplets (LDs) (19). These results suggest that caveolae are involved in lipid accumulation in adipocytes and may promote fatty acid uptake (20). However, the molecular mechanisms of caveolae-dependent fat uptake have remained obscure.

Eps15 homology domain containing protein 2 (EHD2) localizes to the caveolar neck region (6, 21, 22). The protein belongs to the dynamin-related EHD ATPase family, which comprises four members in human (EHD1-4), and shows strong expression in human adipose and muscle tissue (human protein atlas) (23). EHD is built of an N-terminal GTPase (G)-domain, which mediates dimerization and oligomerization, a helical domain containing the membrane binding site, and a C-terminal regulatory Eps15 homology (EH)-domain. The proteins exist in a closed auto-inhibited conformation in solution (24). When recruited to membranes, a series of conformational changes aligns the phospholipid binding sites with the membrane and facilitates oligomerization of EHD2 into ring-like structures (25–27).

Down-regulation of EHD2 in cell culture results in decreased surface association and increased mobility of caveolae, whereas EHD2 overexpression stabilizes caveolae at the plasma membrane (9, 21, 22, 28). This led to the hypothesis that formation of an EHD2-ring at the neck of caveolae restricts caveolar mobility within the membrane. In agreement with this hypothesis, EHD2 assembles in an ATP-dependent fashion into ring-like oligomers *in vitro* and induces the formation of tubular liposomes with an inner diameter of 20 nm, corresponding to the diameter of the caveolar neck (24). Whether EHD2 also controls caveolar membrane dynamics *in vivo* and what the physiological consequences of EHD2 loss at the organismic level are, is unknown.

In this study, we found that EHD2 KO mice display enlarged lipid accumulation in white and brown adipose tissue, and increased lipid droplets (LDs) in caveolae-harboring cell types like adipocytes or fibroblasts. In adipose tissue lacking EHD2, caveolae were frequently detached from the plasma membrane and displayed elevated mobility. Furthermore, in two obesity mouse models, as well as in white adipose tissue of obese patients, reduced EHD2 expression and an increased number of detached caveolae were found in visceral fat. Our data establish EHD2 as a negative regulator of caveolae-dependent lipid uptake and implicate a role of caveolar stability and dynamics for lipid homeostasis and obesity.

## Results

### Generation of EHD2 knockout mice

To examine the physiological function of EHD2, a mouse strain with LoxP recognition sites surrounding exon 3 and intron 3 of the *Ehd2* gene was engineered (Fig. 1A). Exon 3 encodes part of the highly conserved GTPase domain (residues 137-167), and its deletion is predicted to result in nonfunctional protein. Following global removal of exon 3 by crossings with a germ-line specific Cre-deleter strain, offspring mice were back-crossed with the C57BL6/N mouse strain for five generations, yielding a global EHD2 KO mouse model. Genotyping of offspring confirmed the successful deletion of EHD2 exon 3 in EHD2 del/del animals (Fig. S1A) and real-time PCR revealed the absence of EHD2 mRNA in the EHD2 del/del tissue (Fig. 1B). Western blot analysis indicated the complete loss of EHD2 in WAT and various other tissues of EHD2 del/del mice compared to abundant EHD2 expression in EHD2 +/+ mice (Fig. 1C, Fig. S1B-D). Cav1 and Cavin1 protein levels remained grossly unaltered upon loss of EHD2 (Fig. 1C, Fig. S1D). Accordingly, immunostaining of cryostat sections obtained from EHD2 del/del BAT did not reveal apparent differences in Cav1 or Cavin1 expression and localization (Fig. S1E), whereas EHD2 staining was completely abolished (Fig. 1D).

**Fig.1.**
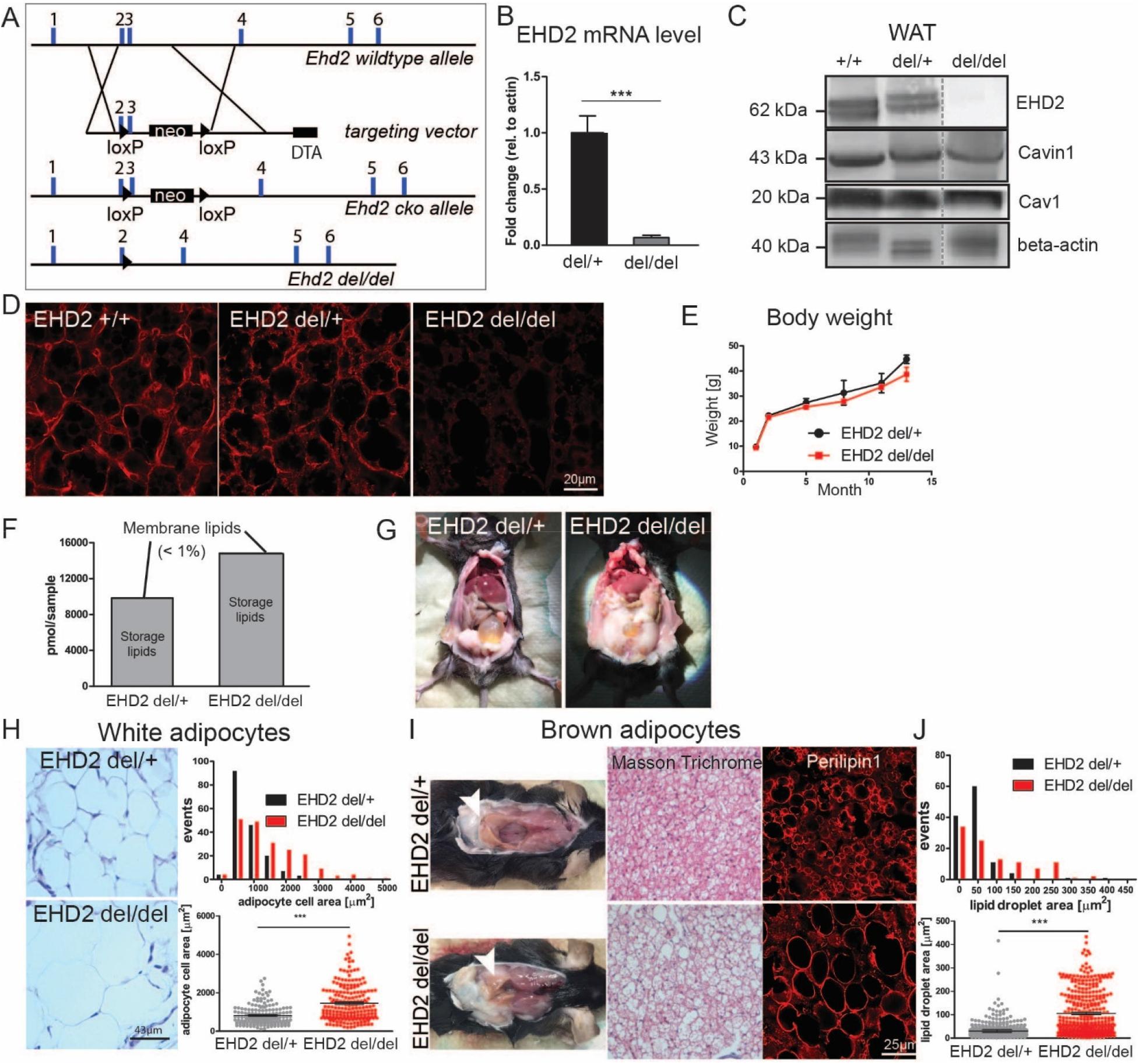
Loss of EHD2 results in increased lipid accumulation in white and brown adipose tissue. **A** Generation of the EHD2 KO mouse model. A targeting vector containing a pGK-Neomycin (neo) cassette and loxP sites flanking exon 3 was placed in the EHD2 wt allele. EHD2 del/del mice were obtained by breeding with Cre-deleter mouse strain (DTA – diphtheria toxin A). **B** EHD2 mRNA level in EHD2 del/+ and EHD2 del/del mice (mRNA from BAT, n = 5). **C** Western Blot analysis of WAT of EHD2+/+, +/− and −/− mice against EHD2, Cav1 and Cavin1. **D** EHD2 immuno-staining in BAT cryostat sections from EHD2 +/+, del/+ and del/del mice. **E** The body weight was monitored over 12 months (n = 7). **F** Lipid composition analysis of 15 μg WAT obtained from EHD2 del/+ or EHD2 del/del mice. **G** EHD2 del/+ and EHD2 del/del mice during preparation. **H** Masson Trichrome staining of WAT paraffin sections of EHD2 del/+ and EHD2 del/del. Detailed analysis of the adipocytes cell size (K, n(del/+) = 172/3, n(del/del) = 199/3). **I** EHD2 del/del mice showed decreased BAT in the neck region. Instead, WAT was integrated into the BAT depots. **J** Masson Trichrome staining of EHD2 del/del BAT paraffin sections and BAT cryostat sections stained against the LD coat protein Perilipin1. LD size was measured in BAT cryostat sections (M, n(del/+) = 118/3, n(del/del) = 104/3). Line graph represents mean +/− SE, column bar graphs show mean + SE, normal distributed groups were analyzed by t-test, not normally distributed values with Mann Withney U test, * P<0.05, *** P<0.0001. For comparison to C57BL6/N, see also Fig. S1–S3.

### Loss of EHD2 results in increased lipid accumulation

EHD2 del/del mice were born in normal Mendelian ratios, were fertile and did not show any obvious phenotype upon initial inspection. In contrast to the reported loss of white fat mass in Cav1 and Cavin1 KO mouse models (4, 16, 17), one year-old EHD2 del/del male mice were not apparently lipodystrophic and did not display any detectable weight difference when compared to EHD2 del/+ mice (Fig. 1E). However, analysis of the lipid composition in WAT indicated an increased amount of storage lipids (mainly triacylglycerol) in EHD2 del/del mice compared to EHD2 del/+ (Fig. 1F). This suggested that these mice suffered from a more subtle phenotype. Indeed, several of the EHD2 del/del mice showed increased deposits of epigonadal and periinguinal white fat (Fig. 1G) indicating that the storage capacity of normal adipose tissue was dysregulated.

When analyzed at the cellular level, white adipocytes of EHD2 del/del mice showed an increased cell size compared to adipocytes from EHD2 del/+ mice or adult C57BL6/N mice (Fig. 1H, Fig. S2A), likely due to increased lipid storage. In contrast to EHD2 del/+ BAT with its distinct brown appearance, EHD2 del/del BAT showed increased beige and white coloring (Fig. 1I). Histological inspections of BAT paraffin and cryostat sections stained against the LD coat protein Perilipin1 indicated an increased LD size in EHD2 del/del BAT compared to EHD2 del/+ or C57BL6/N mice (Fig. 1J, Fig. S2B-C). As no significant differences lipid accumulation (Fig. S2) were found in EHD2 del/+ and C57BL6/N male mice and to reduce animal numbers, the following experiments were carried out with EHD2 del/+ as control group to EHD2 del/del male mice.

Based on the increased lipid accumulation in EHD2-lacking adipocyte tissue, we further investigated if adipocyte differentiation is altered in EHD2 del/del WAT. However, adipogenic marker genes like PPARγ, Retn or Serpina3k displayed no significant expression change in EHD2 del/del WAT compared to EHD2 del/+ WAT (Fig. S2D). The weight adaptation of both genotypes suggested the occurrence of compensatory mechanisms. First, we looked for changes of adiponectin, leptin and insulin plasma levels but did not detect any significant difference in mice lacking EHD2 compared to EHD2 del/+ mice (Fig. S3). Free fatty acid concentration in blood plasma was slightly reduced in EHD2 del/del suggesting a possible increase in fatty acid uptake (Fig. S3). Furthermore, we observed a down-regulation of genes involved in *de novo* lipogenesis in isolated WAT obtained EHD2 del/del vs. EHD2 del/+ mice or in primary adipocyte cell cultures (Fig. S5B-D) suggesting that downregulation of lipogenesis is an active mechanism that partially compensates for the increased lipid accumulation.

### Increased lipid droplet size in adipocytes lacking EHD2

To characterize the mechanisms of increased lipid accumulation in EHD2 del/del fat cells, lipid metabolism was investigated in cultured adipocytes. Primary pre-adipocytes were isolated from WAT of EHD2 del/+ and EHD2 del/del mice and differentiated into mature adipocytes, followed by BODIPY staining to measure LD sizes. Undifferentiated EHD2 del/del pre-adipocytes showed increased LD size, a difference that was even more pronounced in differentiated EHD2 del/del adipocytes compared to EHD2 del/+ (Fig. 2A-B). 3D reconstruction of EHD2 del/del differentiated adipocytes illustrates an extensively increased volume of some LDs (Fig. 2C).

**Fig. 2:**
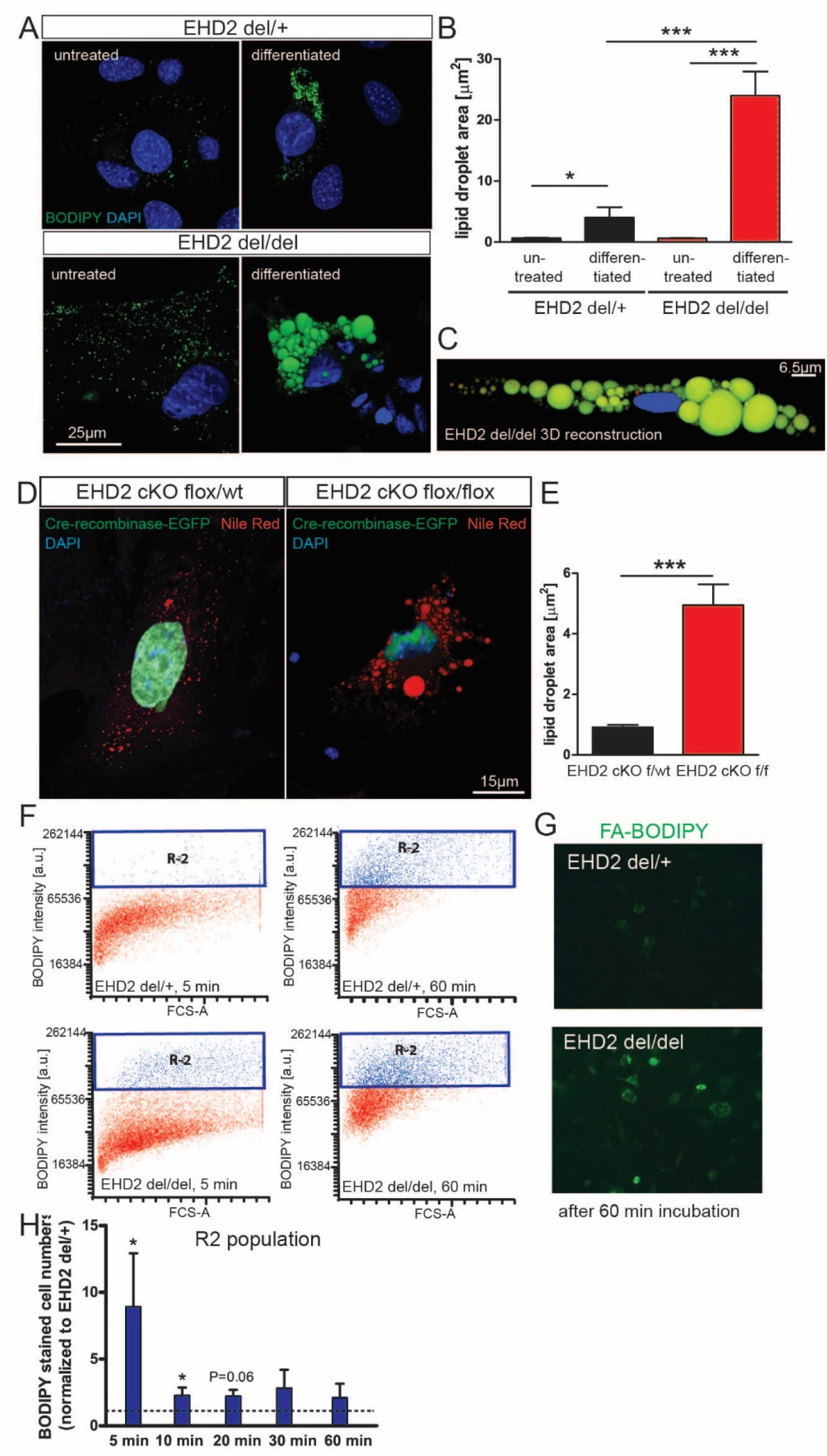
EHD2 del/del adipocytes show faster fatty acid uptake. **A-C** Analysis of LD size in EHD2 del/del and EHD2 del/+ adipocytes by staining with BODIPY (untreated: n(del(+) = 74/3, n(del/del) = 60/3); differentiated: n(del/+) = 132/3, n(del/del) = 95/3). 3D reconstruction of EHD2 del/del differentiated adipocyte (C). Green – LDs, blue – nucleus. **D-E** Cultivated EHD2 cKO flox/wt or flox/flox adipocytes were transfected with Cre recombinase-EGFP to induce EHD2 deletion and differentiated for 5 days, lipid droplets were stained with Nile Red for analyzing (n(flox/wt) = 74/2, n(flox/flox) = 82/2). **F-H** Fatty acid uptake assay in differentiated EHD2 del/+ and EHD2 del/del adipocytes. Dodecanoic acid-BODIPY uptake was measured after 5, 10, 20, 30 or 60 min, and R1 population indicates positively stained cells (illustrated in red in graph F). R2 populations (blue) correspond to higher BODIPY staining intensity in cells and represent adipocytes with increased amount of dodecanoic acid taken up (shown in blue in graphs F, H). Normalization of EHD2 del/del R2 population relative to EHD2 del/+ R2 (H, n(del/+) = 6/3 experiments, n(del/del) = 8/3 experiments). Example images of differentiated adipocytes treated with dodecanoic acid for 60 min (G, scale bar 40 μm). Column bar graphs illustrate mean +/− SE, t-test or Mann Withney U test were used to calculate significance, * P<0.05; *** P<0.0001. See also Fig. S4 and S5.

We addressed the possibility that increased lipid uptake is a secondary effect of EHD2 deletion mediated via putative organ cross-talk. We therefore repeated the experiments with cultivated adipocytes derived from EHD2 cKO flox/flox mice, in which EHD2 expression was down-regulated by expression of Cre recombinase via viral transfection (AAV8). Again, EHD2 removal led to increased LD growth (Fig. 2D-E), indicating a cell-autonomous function of EHD2 in controlling lipid uptake.

LD growth is mainly mediated by extracellular fatty acid uptake and conversion into triglycerides, whereas increased glucose uptake and *de novo* lipogenesis play minor roles in this process (29, 30). Fatty acids and lipids are present at high concentrations in fetal bovine serum (FBS). Addition of delipidated FBS during adipocyte differentiation resulted in complete loss of LD in both genotypes (Fig. S4A), whereas glucose-depletion led to a general impairment of adipocyte differentiation. Importantly, enlarged LDs were still observed in KO adipocytes even under conditions of glucose depletion (Fig. S4A). These data led us to hypothesize that the increased LD size in EHD2 del/del adipocytes may be a consequence of increased fatty acid uptake, in line with the suggested function of caveolae in fatty acid uptake (31–33).

To test this possibility directly, we monitored the uptake of extracellularly added fatty acids into differentiated adipocytes using BODIPY-labelled dodecanoic acid (FA12) paired with FACS analysis. After 5 min, only a minor fraction of EHD2 del/+ adipocytes displayed intense BODIPY staining (R2, for definition see Fig. 2F). This R2 population increased to more than 30% after 60 min of FA12 treatment. EHD2 del/del adipocytes displayed increased BODIPY staining at both early and late time points (Fig. 2F-H; Fig. S4B-D), indicating accelerated lipid uptake. This conclusion was further supported by light microscopy imaging, which revealed more intense BODIPY staining of EHD2 del/del cells compared to EHD2 del/+ adipocytes after 60 min of fatty acid incubation (Fig. 2G). In contrast, EHD2 del/+ and EHD2 del/del adipocytes did not differ with respect to their ability to take up extracellularly added glucose (Fig. S4E-G). Previously, an involvement of EHD2 in the autophagic engulfment of LDs (lipophagy) was suggested (34). However, inducing starvation by incubation of differentiated adipocytes with Hank's balanced salt solution (HBSS) revealed no differences in the release of stored lipids in EHD2 del/+ and EHD2 del/del adipocytes, and both genotypes displayed similar reductions in lipid accumulation (Fig. S5A). These data indicate that loss of EHD2 does not affect the release of fatty acids or lipophagy, but specifically controls LD size by regulating fatty acid uptake.

### Loss of EHD2 results in detachment of caveolae from the plasma membrane *in vivo*

Given the known role of caveolae in the uptake of fatty acids (16), we hypothesized that EHD2 may restrict fatty acid uptake by controlling caveolar function and/ or dynamics. To test whether loss of EHD2 affects caveolar morphology, WAT and BAT were analyzed by electron microscopy (EM). Caveolae in EHD2 del/+ BAT were mostly membrane-bound and displayed the characteristic flask-shaped morphology (Fig. 3A, white arrows, ratio detached/membrane bound caveolae = 0.27). Strikingly, an increased number of caveolae appeared detached from the plasma membrane in BAT isolated from EHD2 del/del mice compared to EHD2 del/+ controls, as judged from the complete closure of the lipid bilayer in the plasma membrane and the caveolae (Fig. 3A, B, black arrows, ratio detached/membrane bound caveolae = 1.75). The total number of caveolae, as well as the caveolar diameter and size, were unchanged in brown adipocytes lacking EHD2.

**Fig. 3:**
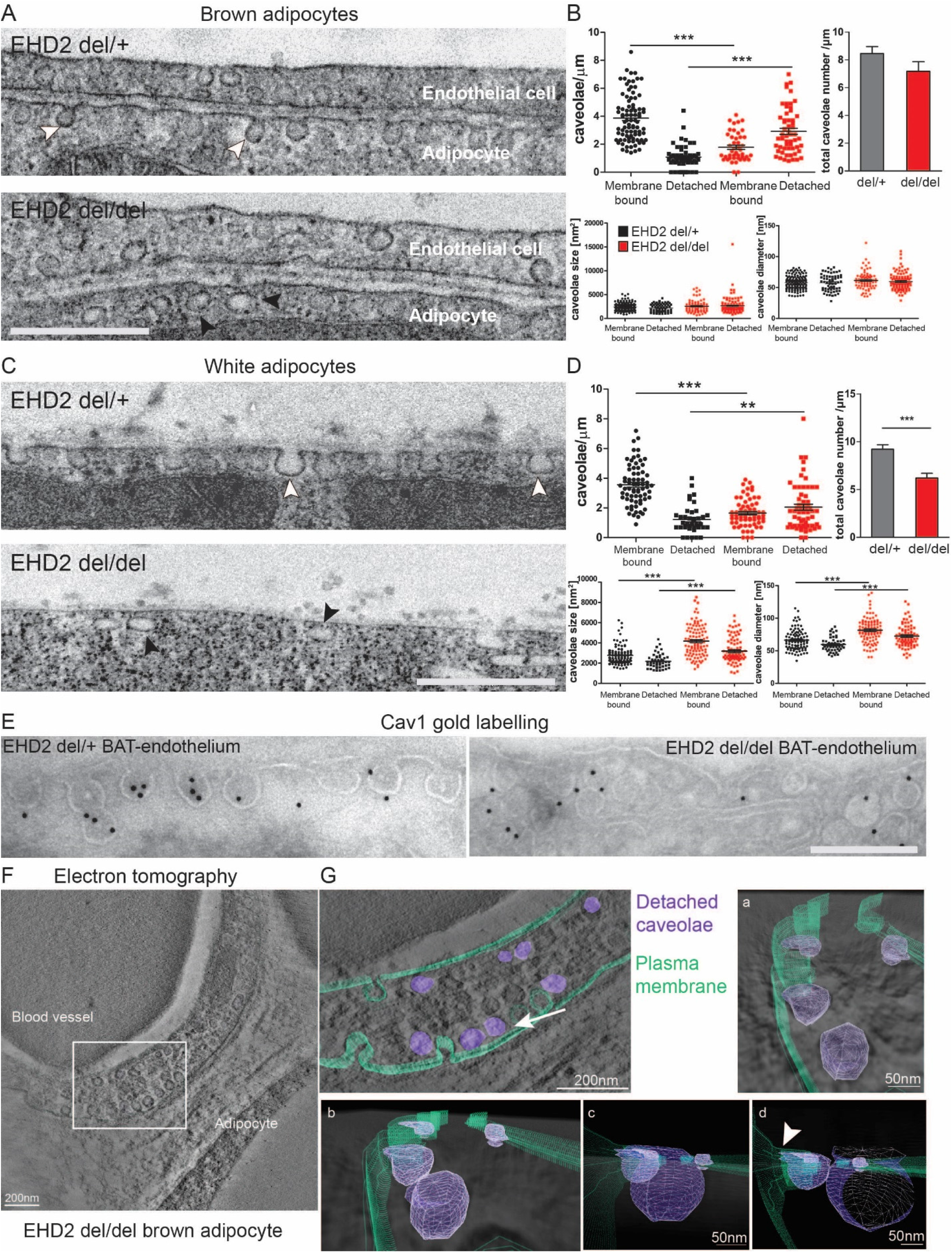
Loss of EHD2 resulted in detached caveolae *in vivo*. **A-B** Representative EM images of BAT from EHD2 del/+ and del/del mice and systematic analysis (caveolae number: n(del/+) = 140/3, n(del/del) = 100/3; caveolae size and diameter: n(del/+) = 201/3, n(del/del) = 171/3). Scale bar 500 nm. **C-D** EM images of EHD2 del/+ and del/del WAT (caveolae number: n(del/+) = 108/3, n(del/del) = 124/3; caveolae size and diameter: n(del/+) = 151/3, n(del/del) = 185/3). Scale bar 500 nm. **E** Representative image for EM gold immunolabeling against Cav1. Control labeling did not reveal specific staining. Scale bar 200 nm. **F-G** Electron tomogram of a 150 nm EHD2 del/del BAT section (F). The 3D model contains the plasma membrane (G, green) and the detached caveolae (violet). Detachment of caveolae was observed by changing the viewing angle (white arrow indicates the direction). Closer inspection of cell membrane and caveolae clearly showed displacement of caveolae from the membrane. The 3D model also revealed attachment of caveolae to the membrane (arrow head). Graphs illustrate each replicate with mean +/− SE, column bar graphs illustrate mean + SE t-test or Mann Withney U test were used to calculate significance, ** P<0.001; *** P<0.0001. See also Movie S1

An increased number of detached caveolae were also observed in EHD2 del/del white adipocytes compared to EHD2 del/+ cells from littermate controls (Fig. 3C, D, black and white arrow heads, ratio detached/membrane bound caveolae (del/del) = 1.2 vs. ratio (del/+) = 0.24). In white adipocytes, the total number of caveolae was reduced, while both caveolar size and diameter were increased in EHD2 del/del compared to EHD2 del/+ animals (Fig. 3D). Cav1 immunogold labeling confirmed that the round vesicles close the plasma membrane, indeed, were detached caveolae (Fig. 3E). 3D visualization of EHD2 del/del brown adipocyte by electron tomography (ET) further indicated that the majority of detached caveolae in 2D EM images were not connected to the plasma membrane (Fig. 3F, G, Movie S1) but localized 20-30 nm underneath (Fig. 3G a, b), although some caveolae close to the plasma membrane showed thin connections (Fig. 3G b, d, white arrow head). Taken together, EM and ET reveal an increased detachment of caveolae from the plasma membrane in EHD2 del/del adipocytes, suggesting a crucial function for EHD2 in the stabilization of caveolae at the plasma membrane.

### Increased caveolar mobility in EHD2 knockout cells

To further dissect the interplay of caveolar mobility and LD growth at the molecular level, we investigated caveolar mobility and endocytosis in mouse embryonic fibroblasts (MEFs) by total internal reflection fluorescence (TIRF) microscopy. MEFs derived from EHD2 +/+ and del/del mice were transfected with pCav1-EGFP to label single caveolae. As illustrated in Fig. 4A, regions of moderate Cav1 expression were investigated to ensure that distinct Cav1 spots were observed during the analysis. Live TIRF imaging of EHD2 +/+ MEFs showed a slow or no continuous movement for the majority of investigated caveolae (Movie S2). However, single caveolae moved along the plasma membrane or left the TIRF illumination zone towards the inside of the cell, indicative of their spontaneous detachment, as previously reported (21). Strikingly, movement and velocity of caveolae was greatly increased in EHD2 del/del MEFs (Movie S3), not allowing Cav1 single spots to be tracked. Line scan analysis revealed a greatly reduced number of fixed, non-moving Cav1 spots (referred to as lines in Fig. 4B) in EHD2 del/del MEFs compared to EHD2 +/+ cells. Moreover, a larger number of highly mobile Cav1 sparks, reflecting fast moving caveolae, was found in EHD2 del/del cells (Fig. 4A, B). Re-expression of EHD2 in EHD2 del/del MEFs reduced the mobility of caveolae, often leading to their immobilization (Fig. S6, Movie S4).

**Fig. 4.**
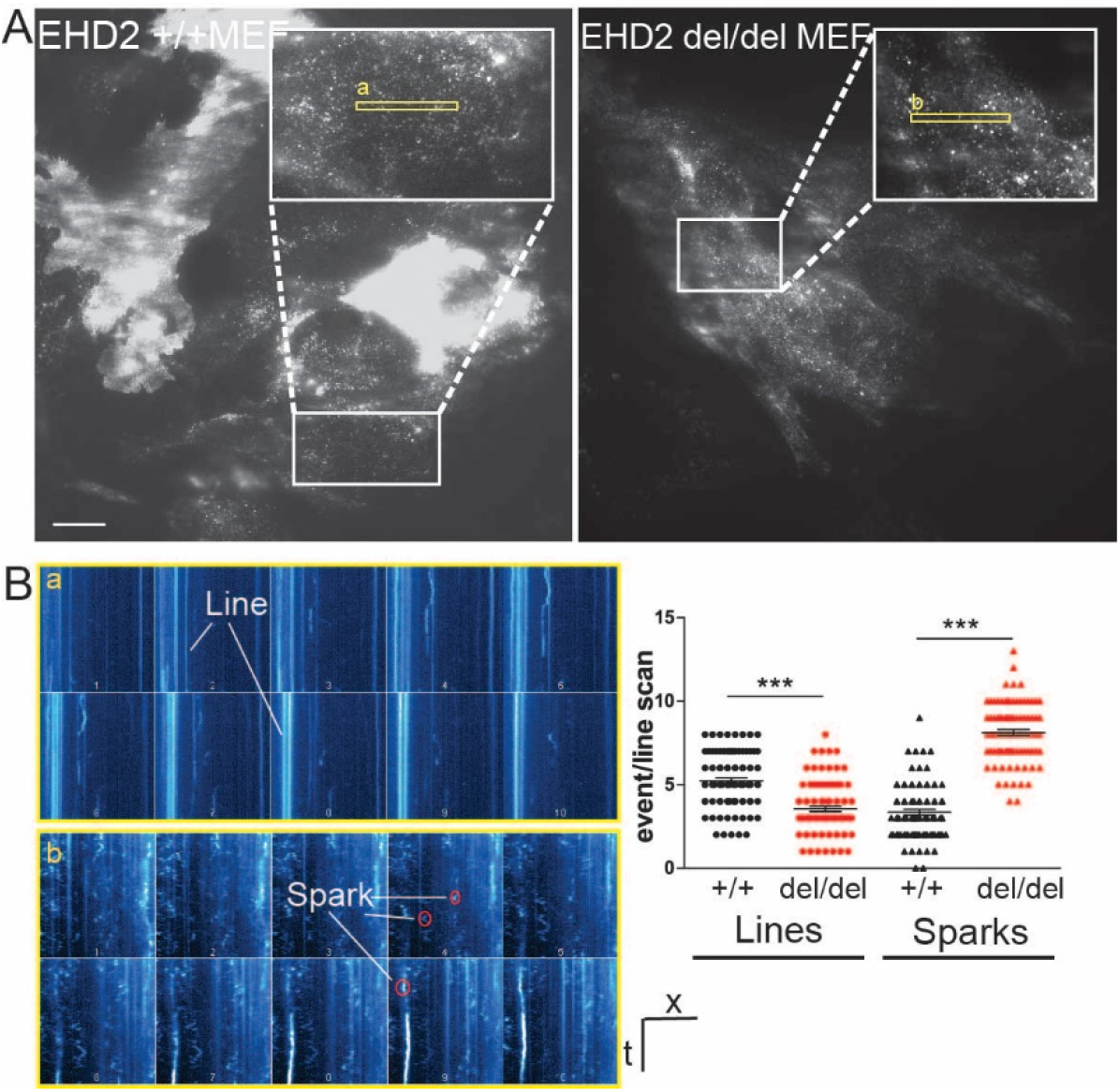
Enhanced caveolar mobility in cells lacking EHD2. **A-B** TIRF live-imaging of EHD2 +/+ and del/del MEFs expressing pCav1-EGFP. Line scan analysis of the recorded Cav1 intensities revealed for fixed, non-moving caveolae lines and for fast moving caveolae single sparks (as illustrated in a and b, n(+/+) = 90/3; n(del/del) = 92/3; each replicate is represented with mean +/− SE), ***P<0.0001. See also Fig. S6 and Movie S2-4.

### Determinants of EHD2-mediated fatty acid uptake

We further characterized the determinants of the observed lipid uptake in MEFs. Similar to EHD2 del/del adipocytes, EHD2 del/del MEFs showed increased lipid accumulation and LD size after adipogenic differentiation, as illustrated by Oil red O staining (Fig. S7A) and BODIPY staining (Fig. S7B, C). Furthermore, both storage and membrane lipids were increased in MEFs lacking EHD2 (Fig. S7D). Re-expression of an EGFP-tagged EHD2 version in in EHD2 +/+ and del/del MEFs rescued the observed LD phenotype, even reducing the size of LDs compared to EGFP expressing cells (Fig. 5A, B). These data indicate a general and cell autonomous role of EHD2 in the control of LD growth and size that is not restricted to fat cells.

**Fig. 5:**
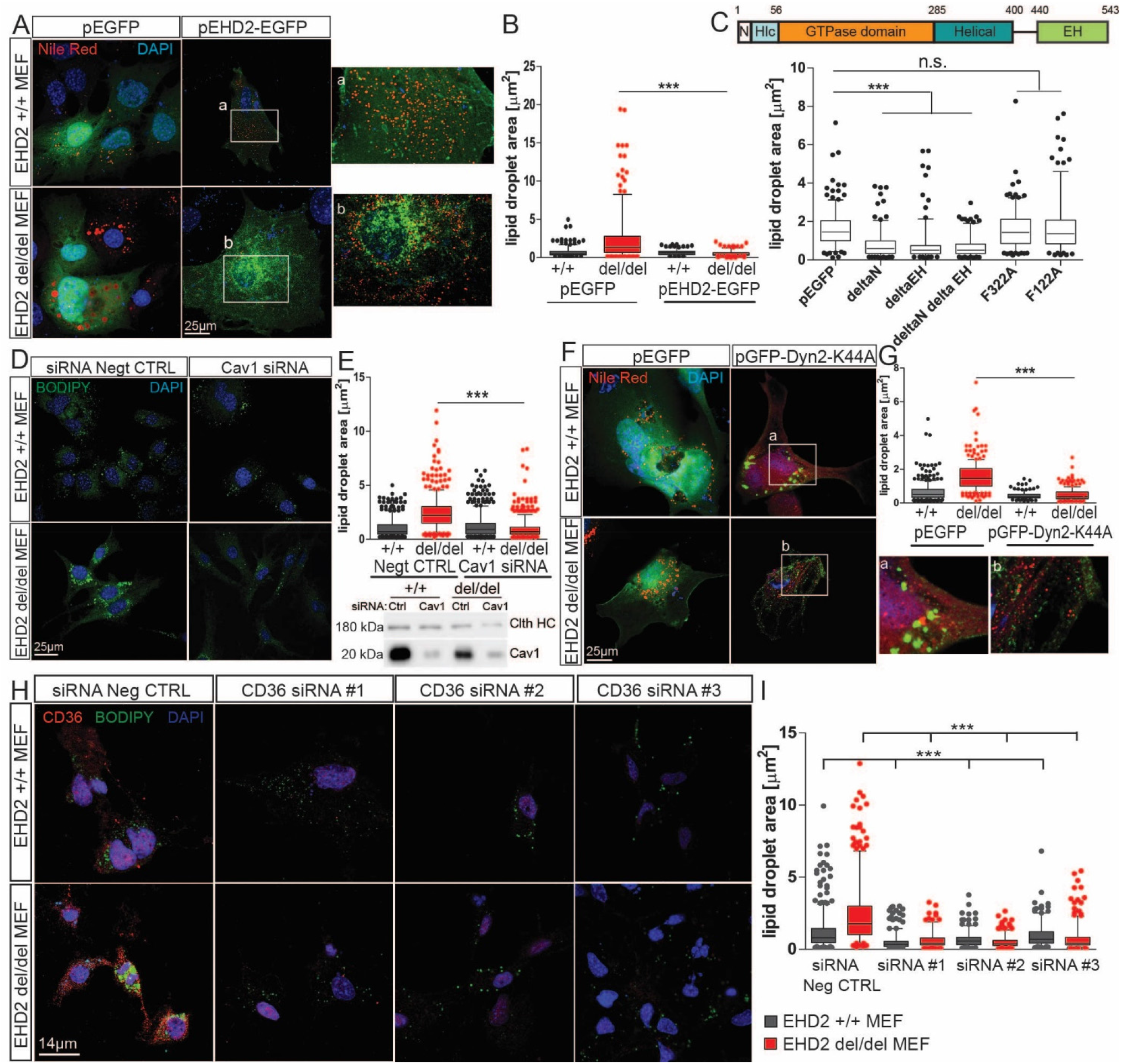
EHD2-mediated fatty acid uptake depends on Cav1, Dyn2 and CD36. **A-B** EHD2 +/+ and del/del MEFs were transfected with either pEGFP or pEHD2-EGFP, incubated for 48 h and afterwards treated for 6 h with oleic acid and Nile Red staining was performed to determine LDs (pEGFP: n(+/+) = 309/3, n(del/del) = 310/3; pEHD2-EGFP: n(+/+) = 218/4, n(del/del) = 184/4). **C** EHD2 constructs were transfected in EHD2 del/del MEFs and after 48h treated with oleic acid and Nile Red staining was performed to determine LDs (n(pEGFP) = 275/3; n(pEHD2-deltaN-EGFP) = 193/3; n(pEHD2-deltaEH-EGFP) = 197/3; n(pEHD2-deltaN-EH-EGFP) = 196/3; n(pEHD2-F322A-EGFP) = 204/3; n(pEHD2-F122A-EGFP) = 212/3). **D-E** EHD2 del/del MEFs were treated with Cav1 siRNA and lipid droplets were stained with BODIPY (negative control: n(+/+) = 504/3, n(del/del) = 530/3; Cav1 siRNA: n(+/+) = 521/3, n(del/del) = 558/3); Clth HC – clathrin heavy chain. **F-G** EHD2 +/+ and del/del MEFs were transfected with either pGFP-Dyn2-K44A, incubated for 18 h and afterwards treated for 6 h with oleic acid and Nile Red staining was performed to determine LDs (pEGFP: n(+/+) = 309/3, n(del/del) = 233/4; pGFP-Dyn2-K44A: n(+/+) = 136/3, n(del/del) = 237/4). **H-I** LD size after CD36 siRNA knockdown in EHD2 +/+ and del/del MEFs (D, negative control: n(+/+) = 584/6, n(del/del) = 475/6; CD36 siRNA#1: n(+/+) = 341/3, n(del/del) = 249/3; CD36 siRNA#2: n(+/+) = 412/3, n(del/del) = 468/3; CD36 siRNA#3: n(+/+) = 251/3, n(del/del) = 368/3; graph illustrates each replicate with mean +/− SE, 2-way ANOVA test were used to calculate significance between siRNA negative CTRL and siRNA, t-test was used between +/+ and del/del data). Box plots indicate mean +/− SE and single replicates of 5% of maximal and minimum values are illustrated, t-test or Mann Withney U test were used to calculate significance, *** P<0.0001. See also Fig. S7.

To identify the functional regions within EHD2 that are crucial for regulating lipid uptake, we transfected EHD2 del/del MEFs with various EGFP-tagged EHD2 deletion constructs. EHD2 constructs lacking the N-terminus, the EH-domain or both, rescued EHD2 loss, resulting in smaller LDs (Fig. 5C, S7E). In contrast, expression of single EHD2 mutants affecting membrane binding (F322A) or oligomerization/ATPase activity of EHD2 (F122A) did not reduce LD size, indicating a crucial role of these properties for EHD2 function.

To analyze if the observed phenotype in EHD2 del/del MEFs was dependent on caveolae, MEFs lacking EHD2 were treated with Cav1 siRNA to eliminate caveolae. LD size in EHD2 del/del MEFs was significantly decreased following depletion of Cav1 (Fig. 5D, E), indicating that the effects of EHD2 on LD size require and likely are mediated by caveolae.

It was previously reported that dynamin is located on caveolae (35). Consistent with a role of dynamin in caveolae function, overexpression of a dynamin 2 dominant negative mutant (pGFP-Dyn2-K44A, Fig. 5F, G) completely abolished the size increase of LDs in EHD2 del/del MEFs, suggesting a role for dynamin in fatty acid uptake via caveolae.

Previous work has implicated the fatty acid binding membrane protein CD36 in caveolae-dependent fatty acid uptake (31, 36, 37). To probe a possible function of CD36 in EHD2-dependent lipid uptake, CD36 expression in MEFs was downregulated by treatment with either one of three specific CD36 siRNA. Antibody staining confirmed the efficient knockdown of CD36 in EHD2 +/+ and del/del MEFs (Fig. 5H). Removal of CD36 in EHD2 +/+ and del/del MEFs dramatically decreased the size of LDs compared to a control siRNA treated cells (Fig. 5H). Hence, the observed enlargement of LDs in cells lacking EHD2 depends on CD36. These converging lines of evidence suggest that caveolae dynamics is key to the regulation of fatty acid internalization and that this is coupled to the previously shown importance of CD36 and caveolin as fatty acid binding proteins.

### Decreased EHD2 expression in genetic obesity models or in diet induced obesity

Our data indicate an EHD2-dependent regulation of lipid uptake in adipose tissue. If this were of physiological relevance, one might expect EHD2 expression to be dysregulated in obese mice and men. We therefore investigated if EHD2 expression is altered in two obesity-related mouse models, ob/ob and NZO (38). Indeed, WAT obtained from ob/ob and NZO mice showed reduced EHD2 expression compared to C57BL6/N mice fed by standard diet (Fig. 6A). When investigating adipocytes from ob/ob mice, a higher proportion of detached caveolae were found in the obesity mouse model (ratio detached/membrane bound caveolae = 1.4 vs. 0.35 in C57BL6/N mice fed with standard diet, Fig. 6B, C). In addition, we analyzed EHD2 expression in visceral and subcutaneous WAT from patients ranking in their body mass index (BMI) from normal to morbid obesity (BMI<25 – BMI >40, Fig. 6D, E). In both depots, EHD2 expression was highest in normal weight subjects and was significantly lower in overweight and obese people, whereas EHD2 expression did not differ between different obesity stages. These data imply that EHD2 expression is regulated by lipid uptake and load and suggest that EHD2-mediated caveolar dynamics may be altered in obesity.

**Fig. 6:**
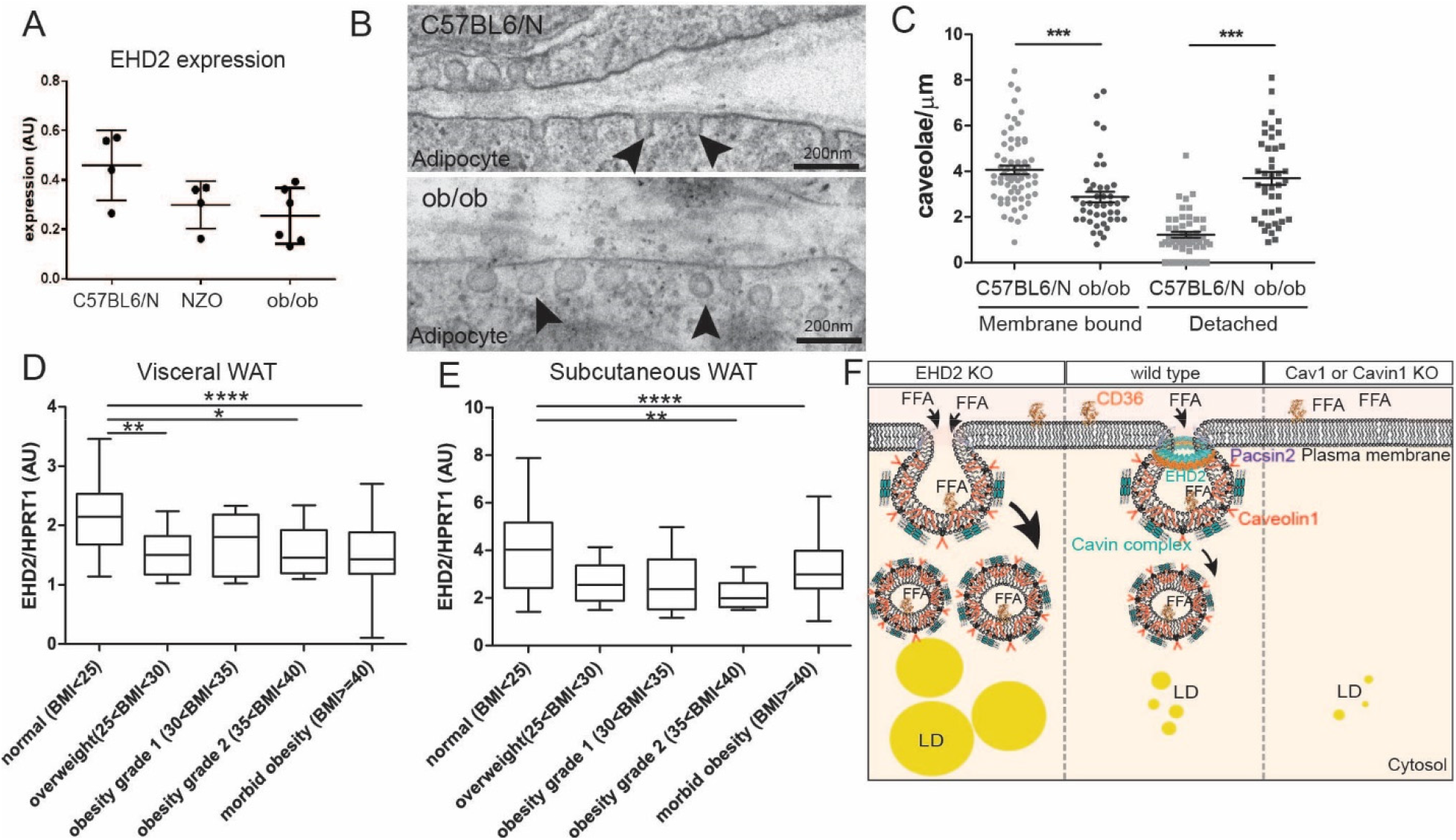
Decreased EHD2 expression in diet induced obesity. **A** EHD2 expression level was analyzed in fat tissue of ob/ob or NZO mouse models compared to C57BL6/N mice (n = 5). **B-C** Investigation of caveolae by EM imaging (n(ob/ob mice) = 85/2; n(C57BL6/N) = 117/2;). **D-E** EHD2 expression in visceral (E) or subcutaneous WAT (F) in obese patients (n(normal) = 31; n(overweight) = 23; n(obesity grade 1) = 7; n(obesity grade 2) = 17; n(obesity grade 3) = 202). **F** The model illustrates how fatty acid uptake is affected by caveolae function. In the absence of caveolae, lipid uptake is reduced resulting in smaller lipid droplets compared to normal conditions. In the absence of EHD2, lipid uptake is increased, suggesting a regulatory function of EHD2 in caveolae-dependent lipid uptake. FFA – Free fatty acid, LD – lipid droplet. Box plots indicate median with whiskers from maximal to minimum value, or each replicate with mean +/− SE is represented, normal distributed groups were analyzed by t-test, not normal distributed values with Mann Withney U test, * P<0.05, *** P<0.0001.

## Discussion

Here, we identify EHD2 as a negative regulator of caveolae-dependent lipid uptake. Loss of EHD2 resulted in increased lipid accumulation, which was observed in adipose tissue of the whole organism as well as in cell culture-based experiments. Loss of EHD2 was associated with the detachment of caveolae from the plasma membrane, higher caveolar mobility and increased lipid uptake. We demonstrate that caveolae, dynamin 2 and the fatty acid translocase CD36 play a role in the EHD2-dependent lipid uptake pathway. In addition, obese mouse models exhibit decreased EHD2 expression in WAT and in addition, also obese patients showed a reduction in EHD2 gene expression. Thus, our study reveals a cell-autonomous caveolae dependent lipid uptake route that is controlled by EHD2 and modified by metabolic conditions.

For some time, caveolae have been implicated in lipid uptake. Thus, mice lacking caveolae showed reduced fat mass and did not develop any form of obesity. In addition, Pohl et al. (39) observed decreased oleate uptake after expression of a dominant-negative Cav1 mutant. The EHD2 KO mouse model, described here, revealed the opposite phenotype, e.g. a caveolae gain-of-function *in vivo* model. In EHD2 KO mice, caveolae were more often detached from the plasma membrane and showed a higher mobility and faster lipid uptake, resulting in enlarged LDs. Unlike in Cav1 over-expressing mice (19), which also show increased fatty acid uptake, the number of caveolae was not increased in EHD2 KO mice. This supports a model in which not only caveolae number, but also caveolar dynamics play a crucial role in this process (Fig. 6F). Furthermore, such idea is in line with our structural findings that EHD2 can form ring like oligomers that may stabilize the neck of caveolae (24–27), thereby restricting caveolar mobility (21, 22). Based on studies in EHD2 KO NIH 3T3 cell line, Yeow et al. (40) suggested that EHD1 and/or EHD4 could rescue loss of EHD2 during its role in membrane protection. However, we did not find rescue of caveolae detachment and lipid uptake in EHD2 KO cells by other EHD family members.

The detailed pathway of caveolae-dependent cellular lipid uptake has been intensively studied (4, 20, 31, 32, 41), but the exact molecular mechanisms are still unclear. Our observation indicate that fatty acid uptake depends on caveolar dynamics, detachment and most likely caveolae endocytosis and is regulated by EHD2. However, it remains elusive how caveolar shuttling is linked to the growth of lipid droplets. The EHD2 knockout model may be a useful tool to further dissect the molecular lipid uptake mechanism.

The loss of EHD2 on the cellular level led to increased fat deposits on the organismic level, which was particularly evident in older animals. The observed phenotype based on the global loss of EHD2 could be influenced by organ-organ interactions (42). However, we did not find any evidence for differences in adipocyte derived secretory factors, such as leptin (Fig. S3). Furthermore, increased lipid uptake was dependent on caveolae, as shown by Cav1 knockdown experiments, and lipid droplet growth could specifically be induced by viral transfection of Cre recombinase in EHD2 cKO flox/flox, but not in flox/wt adipocytes. Thus, increased lipid uptake is caused by a cell-autonomous, caveolae-dependent mechanism. As the WAT distribution and lipid accumulation is known to differ in male and female mice, further studies are required to analyze potential sex-specific differences in EHD2-dependent lipid uptake.

Despite the observed lipid accumulation in mice lacking EHD2, the body weight of EHD2 del/del and EHD2 del/+ mice was unaltered, suggesting metabolic compensation. In line with this idea, EHD2 del/del WAT and cultivated EHD2 del/del adipocytes showed a significant reduction in expression levels of genes involved in *de novo* lipogenesis like SREBP1 or FAS indicating a strong downregulation of glucose-dependent fatty acid production in fat cells. Similar compensatory mechanisms were noted in patients suffering from obesity (43–45). Remarkably, the expression levels of EHD2 in WAT from obese patients as well as in WAT of two obese mouse models (ob/ob and NZO mice) and of EHD2 del/+ mice treated with a long-term high fat diet were significantly reduced. Thus, expression of EHD2 appears to negatively correlate with adipocyte size, therefore reflecting the situation in the EHD2 KO mouse (see also (46)). We speculate that an imbalance in number, life-time and mobility of caveolae may accompany and possibly actively contribute to the development and progression of obesity. Accordingly, pharmacological approaches to enhance EHD2 expression or its stabilization at the plasma membrane could reduce lipid uptake and consequently help to treat obesity in patients.

In conclusion, our study reveals that EHD2 controls a caveolae-dependent cellular lipid uptake pathway.

## Methods

Please see SI for detailed description of all methods and material.

### EHD2 delta E3 mouse strain generation

The EHD2 targeting construct was generated by insertion of two lox P sequences flanking exon 3 of EHD2 genomic DNA by homologous recombination in *E.coli* as previously described (47) including a pGK Neomycin and a diphtheria toxin A (DTA) cassette. Electroporation of the linearized targeting vector in R1 ES cells was performed. Mice carrying a loxP-flanked Exon 3 of EHD2 gene were mated to Cre deleter mice to generate EHD2 mutant (del/del) mice. After backcrossing the EHD2 del/del mice with C57BL6/N (Charles River, between 20-30 weeks, male) for 6 generations only male EHD2 del/del or EHD2 del/+ (as control) mice were used and littermates were randomly assigned to experimental groups. All animals were handled accordingly to governmental animal welfare guidelines and were housed under standard conditions.

### Obesity Mouse Models

Male NZO/HIBomDife (German Institute of Human Nutrition, Nuthetal, Germany), C57BL/6J (Charles River Laboratories, Sulzfeld, Germany) and B6.V-Lepob/ob/JBomTac (B6-ob/ob) mice (Charles River Laboratories, Calco, Italy) were housed under standard conditions (conventional germ status, 22 °C with 12 hour /dark cycling). NZO and C57BL/6J mice were fed were fed standard chow diet (Ssniff, Soest). Starting at 5 weeks of age B6-ob/ob received carbohydrate free diet (48). Mice were sacrificed at an age of 20-22 weeks.

### Oil Red O staining

LDs in tissue sections or cultivated adipocytes and MEFs were stained with Oil Red O as published by (49).

### Human EHD2 expression

All participants were recruited by University Leipzig (approval numbers: 265-8, 159-12-21052012, and 017-12-23012012) and samples were treated as previously described (50).

### Immunocytostaining and LD staining of cultivated cells

Adipocytes or MEFs were seeded on fibronectin (Sigma) coated glass dishes (ThermoFisher). Cells were washed with PBS, treated with 4% PFA for 10 min and blocking buffer (1%donkey serum/1% TritonX100/PBS) for 20 min. The first antibody was incubated for 1 h, followed by secondary antibody and DAPI stain. For LD staining, BODIPY (Invitrogen, saturated solution) or Nile Red (Sigma, saturated solution) was diluted to 1:1000 in PBS and applied for 30 min. The stained cells were washed and the glass dishes were placed on conventional microscope slides and embedded in ImmoMount. Zeiss LSM700 or Zeiss LSM880 microscopes and ImageJ/Fij were used.

### Transmission Electron microscopy (TEM)

Mice were fixed by perfusion with 4% (w/v) formaldehyde in 0. 1 M phosphate buffer. Tissue blocs were postfixed in phosphate buffered 2.5% (v/v) glutaraldehyde, treated with 1% (v/v) osmium tetroxide, dehydrated in a graded series of ethanol and embedded in the PolyBed^®^ 812 resin (Polysciences Europe GmbH). Ultrathin sections (60-80 nm) were cut (Leica microsystems) and stained with uranyl acetate and lead citrate before image acquisition. Samples were examined at 80 kV with a Zeiss EM 910 electron microscope. Acquisition was done with a Quemesa CDD camera and the iTEM software (Emsis GmbH).

### Electron tomography (ET)

To obtain electron tomograms 250 nm slices of EHD2 del/del BAT were prepared of samples embedded in resin and treated as described for TEM. The samples were tilted from 60 to −60° in 2° steps and examined at 120 kV with a FEI Talos electron microscope. FEI tomography software was used for acquisition of tomograms, detailed analysis and reconstruction was done with Inspect3D, Amira (both obtained from FEI) and IMOD (University of Colorado, USA).

### Fatty acid uptake assay

EHD2 del/+ and EHD2 del/del pre-adipocytes were seeded in 6-well plates (100.000 cells/well) and differentiated in mature adipocytes as described above. The fatty acid uptake assay was performed as described elsewhere (51). Briefly, differentiated adipocytes were starved for 1 h with serum-free DMEM. Next, 2 μM dodecanoic acid (FA12) labelled with BODIPY (Molecular probes #D3822) diluted in serum-free DMEM + 10 μg/ml insulin was added to the adipocytes and incubated for 5-60 min at 37°C followed by FACS. Glucose uptake was measured with 200 μM 2-NBDG (2-deoxy-2-[(7-nitro-2,1,3-benzoxadiazol-4-yl)amino]-D-glucose, molecular probes #N13195) diluted in serum-free DMEM + 10 μg/ml insulin.

## Acknowledgments

We thank Petra Stallerow for taking care of the EHD2 delta E3 mouse strain, Karin Jacobi for advice with the animal application, Vivian Schulz and Carola Bernert for helping with the genotyping of the mice, Claudio Shah for purifying the EHD2 antiserum, and the advanced light imaging facility at MDC for technical support. The authors acknowledge financial support from the Deutsche Forschungsgemeinschaft (DFG; SFB 958/A7 to V.H., A12 to O.D., A13 to A.S.) and support by The Initiative and Networking Fund of the Helmholtz Association.

## Author Contributions

C. M. planned, performed and analyzed all experiments if not otherwise indicated. C.M. and O.D. wrote the manuscript, with input from all authors. S.K. performed and analyzed all EM imaging, C.M. analyzed EM images, S.K. and C.M. performed and analyzed ET. I.L. generated the EHD2 KO mouse model and performed in situ hybridization. W.J. analyzed blood plasma markers and EHD2 expression in obesity mouse models. M.L. helped during TIRF imaging and discussed experiments, A.M. isolated primary MEF and performed the EHD2 Western Blot. E.L. performed lipid droplet staining experiments after Cav1 knockdown. M.K. and M.B. recruited obese patients and handled human samples. A.S., V.H., R.L., C.B. and D. N.M. discussed potential experiments and the manuscript. O.D. wrote the mouse animal application with help of D.N.M. and C.M.

## Supporting Information

### Extended Material and Methods

#### Mouse embryonic fibroblast isolation and immortalization

All animals were handled accordingly to governmental animal welfare guidelines. MEFs were obtained from E14.5 EHD2 +/+ or del/del embryos. Therefore female, pregnant EHD2 del/+ were sacrificed by cervical dislocation, the embryos were dissected and removed from the yolk sac in sterile, cold PBS. For genotyping, a small piece of each mouse embryo tail was harvested followed by complete dissection of the whole embryo. Afterwards, the embryo pieces were treated with 0.25% trypsin/EDTA (Sigma) over night at 4 °C. After aspiration of the trypsin solution, 10 ml culture medium (DMEM/10%FBS/5% penicillin/streptomycin) was added and tissue pieces were break up by pipetting. The cell suspension was transferred in 75 cm^2^ culture flask for cultivation at 37 °C and 5% CO2. Immortalization of isolated primary MEFs was assured by frequently splitting. From passage 15 an increased growth rate was observed suggesting immortalized MEFs. For all experiments MEFs between passage 12 and 32 was used. For LD growth, MEFs were either treated with 10 μg/ml insulin, 2.5 mM dexamethasone, 50 mM IBMX and 25 mM rosiglitazone (all obtained from Sigma) diluted in culture medium (differentiation medium) or 0.016 M oleic acid and 10 μg/ml insulin diluted in DMEM.

#### Primary adipocyte cell culture

Male EHD2 del/+ and EHD2 del/del mice or EHD2 cKO flox/wt or flox/flox were sacrificed by cervical dislocation and gonadal WAT was removed. Adipocytes and stromal vascular fraction (SVF) were isolated after washing the tissue in sterile PBS and digestion by collagenase type II (Sigma C6885). Mature adipocytes floating in the upper phase were transferred in a new flask and diluted with culture medium (DMEM/10%FBS/5% penicillin/streptomycin), SVF was obtained after 5 min centrifugation at 1,000 rpm. After complete tissue break up the adipocyte cell suspension was passed through a 270 μm cell strainer and the cells were plated in 75 cm^2^ culture flask at 37 °C and 5% CO2 whereby pre-adipocytes adhere to the flask and mature adipocytes float in the medium. SVF suspension was cleaned by passing through 70 μm cell strainer. The following day the culture medium was exchanged to remove dead or non-adherent cells. After 5 days, both pre-adipocytes and SVF were split by 0.25% trypsin/EDTA solution and merged for further cultivation. Differentiation to mature adipocytes was induced by 10 μg/ml insulin, 2.5 mM dexamethasone, 50 mM IBMX and 25 mM rosiglitazone diluted in culture medium. If not otherwise mentioned the primary pre-adipocytes were incubated for 5 days with differentiation medium and medium was changed after 2 days. Delipidation of FBS was carried out as described by (1). EHD2 cKO adipocytes were transfected with Cre recombinase-EGFP by using adeno-associated virus particles 8 (AAV8) produced from pAAV.CMV.HI.eGFP-Cre.WPRE.SV40 (addgene, #105545). The adipocyte cell culture was transfected and differentiated for 5 days.

#### Oil Red O staining

LDs in tissue sections or cultivated adipocytes and MEFs were stained with Oil Red O (Sigma #O0625) as published by (2). Briefly, freshly dissected liver of muscle pieces were frozen in liquid nitrogen, embedded in TissueTek and 10 μm cryostat sections (Leica) were prepared. After fixation with 4% para-formaldehyde (PFA, Merck) freshly prepared Oil Red O staining solution was applied for 10 min. The sections were washed with PBS and embedded in ImmoMount (Invitrogen). Cultivated cells were fixed, treated with 60% isopropanol (Merck) for 2 min and then incubated with Oil Red O staining solution for 5 min. After washing with water until complete removal of Oil Red O the stained cells or sections were analyzed by Zeiss Axiovert microscope (20x Zeiss objective). Staining intensity was measured with ImageJ.

#### Histology

EHD2 del/+ and EHD2 del/del mice were anesthetized with 2% ketamine/10%rompun, perfused first by 30 ml PBS and next by 50 ml 4% PFA and tissues were dissected. After 24 h of fixation in 4% PFA, tissues were dehydrated in 3 steps (each 24h) from 70-100% EtOH and afterwards incubated in xylol (Merck) for 48 h. Next, the tissues were embedding in liquid paraffin at ca. 65°C and cooled down on ice. 4 μm paraffin sections were obtained, de-paraffinized and hydrated and Masson Trichrome staining (Kit, Sigma) was applied. Briefly, sections were stained with Bouin solution for 15 min at 60°C, followed by Haematoxylin Gill No. 2 staining for 5 min and incubation in Biebrich-Scarlet-Acid Fuchsin for 5 min. Next, the tissue sections were treated with Phosphotungstic/Phosphomolybdic Acid Solution and Aniline Blue solution both for 5 min, and acetic acid treatment (1%) for 2 min. After extensive washing the sections were dehydrated, incubated in xylol and embedded with Roti Histo Kit (Carl Roth). Images were obtained at Zeiss Axiovert100 microscope.

#### Immunohistostaining of cryostat sections

Perfused and fixated EHD2 del/+ and EHD2 del/del mice (as described before) were dissected and the investigated tissue pieces were further fixed for 1-4 h in 4% PFA, transferred to 15% sucrose (in PBS, Merck) for 4 h and finally incubated overnight in 30% sucrose. After embedding in TissueTek, the tissue is frozen at −80 °C. 5-15 μm sections were obtained in a cryostat at – 20 – −30 °C and stored at −20 °C. For immunostainings, the cryostat sections were incubated with blocking buffer (1% donkey serum/1% TritonX100/PBS) for 1 h at room temperature, and treated overnight at 4°C with the first antibody diluted in blocking buffer. After washing with PBS/1%Tween, the secondary antibody was applied for 2 h at room temperature. After completion of the staining, the sections were washed carefully and embedded in ImmoMount. The stained sections were analyzed with Zeiss LSM700 microscope provided with Zeiss objectives 5, 10, 20, 40 and 63x. The obtained images were further investigated by ZEN software and ImageJ/Fij.

#### Transfection and siRNA knockdown

Cultivated MEFs were transfected with the following plasmids pEHD2-EGFP, pEHD2-deltaN-EGFP, pEHD2-deltaEH-EGFP, pEHD2-deltaN-EH-EGFP, pEHD2-F322A-EGFP, pEHD2-F122A-EGFP, pCav1-EGFP (provided from R.L.), pGFP-Dyn2-K44A (provided from M.L., addgene #22301) or pEGFP by lipofectamine 3000 (Invitrogen) accordingly to the manufacture's protocol. Transfected cells were incubated for 48 h and afterwards the treated cells were analyzed by confocal microscopy or TIRF. siRNA knockdown of CD36 or Cav1 was performed in freshly split MEFs by electroporation with the GenePulser XCell (Biorad). Briefly, MEFs were split as described before and the obtained cell pellet was resuspended in OptiMEM (Gibco). After cell counting, the MEF cell suspension was diluted to 1.5×10^6^ cells/ml and 300 μl were transferred into electroporation cuvettes (2 mm, Biorad). CD36 or Cav1 stealth siRNA and siRNA negative control (medium GC content, CD36 siRNA#1 GGAAUUUGUCCUAUUGGCCAAGCUA, CD36 siRNA#2 CCAAGUCUUCUAUGUUCCAAACAAG; CD36 siRNA#3 CCAAUAACUGUACAUCUUAUGGUGC) was added to a final concentration of 200 nM. After carful mixing, the cuvettes were placed into the electroporation device and the pulse (160 μOHM, 500 μF, ∞ resistance) was applied. The electroporated cells were cultivated in DMEM/10%FBS for 48 h before the experiments were started. Successful siRNA knockdown was monitored by CD36 antibody staining.

#### TIRF live imaging of caveolae movement

MEFs transfected with pCav1-EGFP were incubated for 48 h on fibronectin coated cover slips (25 mm diameter). Samples were mounted in Attofluor Cell Chamber (Thermo) in a physiological buffer (130 mM NaCl, 4 mM KCl, 1.25 mM NaH2PO4-H2O, 25 mM NaHCO3, 10 mM glucose, 2 mM CaCl2, 1 mM MgCl2, pH 7.3, 305-315 mOsm/kg).

TIRF imaging was performed on an inverted Microscope (Nikon Eclipse Ti) equipped with a 488 laser (Toptica), an dicroic mirror (AHF, zt405/488/561/640 rpc), a 60x TIRF objective (Nikon, Apo TIRF NA 1.49), an appropriate emission filter (AHF, 400-410/488/561/631-640) and a sCMOS camera (mNeo, Andor). All components were operated by open-source ImageJ-based micromanager software. All experiments were performed at 37 °C. To investigate the movement of single caveolae transfected cells were selected in which regions of individual Cav1 spots were observed (ROIs illustrated in Fig. 4A, enhanced images). Recordings were obtained with the following imaging settings: image size 1776×1760 pixel, 1×1 binning, 500 frames, 200 ms exposure time/frame. For data analysis only the first 150 frames were investigated. After cropping to the specific ROI, kymograph analysis of several positions within the ROIs were carried out using the Reslice function of ImageJ/Fij. Carefully investigation of the kymographs revealed a single, straight line for fixed, not moving caveolae and sparks or short lines for fast moving caveolae.

#### Transmission Electron microscopy (TEM)

Mice were fixed by perfusion with 4% (w/v) formaldehyde in 0. 1 M phosphate buffer and tissues were dissected to 1-2 mm^3^ cubes. For morphological analysis, tissue blocs were postfixed in phosphate buffered 2.5% (v/v) glutaraldehyde. Samples were treated with 1% (v/v) osmium tetroxide, dehydrated in a graded series of ethanol and embedded in the PolyBed^®^ 812 resin (Polysciences Europe GmbH). Ultrathin sections (60-80 nm) were cut (Leica microsystems) and stained with uranyl acetate and lead citrate before image acquisition. For immuno-labeling, samples were fixed by perfusion as described above, but postfixed in phosphate buffered 4% (w/v) formaldehyde with 0.5% (v/v) glutaraldehyde for 1 hour. Samples were further processed as described in Slot and Geuze (Nature protocols, 2007). Briefly, samples were infiltrated with 2.3 M sucrose, frozen in liquid nitrogen and sectioned at cryo temperatures. Sections were blocked and washed in PBS supplemented with 1% BSA and 0.1% glycine. Labeling was performed with an anti-caveolin-1 antibody 1:500 (abcam #2910) and 12 nm colloidal gold (Dianova). Sections were contrasted with 3% tungstosilicic acid hydrate (w/v) in 2.8% polyvinyl alcohol (w/v) (3). Samples were examined at 80 kV with a Zeiss EM 910 electron microscope (Zeiss). Acquisition was done with a Quemesa CDD camera and the iTEM software (Emsis GmbH).

#### *In situ* hybridization

Digoxigenin-labeled riboprobes were generated using a DIG-RNA labeling kit (Roche). In situ hybridizations were performed on 14 μm cryosections prepared from E18.5 wt embryos as previously described (4). To generate an Ehd2 specific in situ probe, a 400 bp fragment was amplified from wildtype cDNA using PCR and primer listed below. The PCR product was cloned into pGEM-Teasy (Promega). T7 and sp6 polymerases were used to generate Ehd2-sense and antisense probes, respectively. EDH2_ISH_FWD: 5’-CAGGTCCTGGAGAGCATCAGC-3’ EDH2_ISH_REV: 5’-GAGGTCCTGTTCCTCCAGCTCG-3’

#### Western Blot

EHD2 protein level in different tissues was examined by Western Blot. Therefore EHD2 +/+, EHD2 del/+ and EHD2 del/del mice were sacrificed by cervical dislocation and organs were dissected and snap frozen in liquid nitrogen. After homogenization of the tissue in 1x RIPA buffer (Abcam) with a glass homogenizer, the tissue lysate was incubated for 1 h on ice followed by 15 min centrifugation at 15,000 rpm. Supernatant was transferred in a fresh tube and protein concentration was measured by NanoDrop. At least 10 μg protein/lane was applied to 4-12% SDS-PAGE NuPage (Invitrogen) and SDS-PAGE was performed accordingly to the manufacture's protocol. Afterwards, proteins were blotted on nitrocellulose membrane (Amersham) at 80 V for 1 h, followed by blocking of the membranes with 5% milk powder (in TBST, 150 mM NaCl, 20 mM Tris-HCl, pH 7.5, 0.1% Tween20) for 2 h at room temperature. To detect EHD2 protein level rabbit-anti-EHD2 (1:2,000) was applied over night at 4 °C. After washing with TBST the secondary antibody goat-anti-rabbit-HRP was added to the membrane for 2 h at room temperature. Detection of EHD2 bands results from ECL detection solution and intensities were obtained by ChemiDoc XRS (Biorad).

#### Antibodies

Anti-EHD2-Rb (self-made), anti-Cav1-Mouse (BD Biosciences #610407), anti-Rabbit IgG HRP (dianova), anti-Mouse IgG HRP (dianova), anti-Perilipin1-Rb (Cell signaling #9349), anti-CD36-Rb (Novus Bio #NB400-144 and abcam #ab133625), anti-Cavin1-Rb (Abcam, #ab48824), anti-Rb-Cy3 (dianova), antimouse IgG-Alexa488 (ThermoScientific #R37114), DAPI (Sigma D9542)

#### Blood plasma analysis

To measure distinct blood plasma parameter related to metabolic changes like adiponectin, insulin or free fatty acids blood was taken from EHD2 del/+ and EHD2 del/del mice immediately after cervical dislocation. All blood samples were taken at 10.00 am. Briefly, mice were opened and the thorax was partly removed to get access to the left heart ventricle, a cannula was inserted and blood samples were taken. After short centrifugation at high speed, the plasma fraction was transferred to a fresh tube and snap frozen in liquid nitrogen. The following assays were used to measure the described blood plasma markers: Plasma insulin levels were measured by Mouse Ultrasensitive Insulin ELISA (80-INSMSU-E10, Alpco). Plasma adiponectin and leptin levels were measured by Mouse Adiponectin/Acrp30 (DY1119) and Mouse/Rat Leptin (MOB00) ELISA kits (R&D Systems). Plasma lipids were quantified with commercially available kits: cholesterol (Cholesterol liquicolour colorimetric assay, Human, Wiesbaden, Germany), triglycerides/glycerol (Triglyceride/Glycerol Calorimetric Assay, Sigma) and non-esterified fatty acids (Wako Chemicals). All measurements were done according to manufacturers' recommendations.

#### Fatty acid uptake assay

EHD2 del/+ and EHD2 del/del pre-adipocytes were seeded in 6-well plates (100.000 cells/well) and differentiated in mature adipocytes as described above. The fatty acid uptake assay was performed as described elsewhere (5). Briefly, after 5 days of differentiation, adipocytes were starved for 1 h with serum-free DMEM. Next, 2 μM dodecanoic acid (FA12) labelled with BODIPY (Molecular probes #D3822) diluted in serum-free DMEM + 10 μg/ml insulin was added to the adipocytes and incubated for 5, 10, 20, 30 and 60 min at 37 °C. After washing twice with ice-cold PBS, 150 μl 0.25% trypsin/EDTA/PBS was applied to detach the cells. The adipocytes were treated with 500 μl ice-cold FACS buffer (HBSS/10%FBS/10 mM EDTA) and the cell solution was transferred to FACS tubes. Shortly before measurement, 1 μl/ml propidium iodide was added. FACS experiments were performed at LSR Fortessa 5Laser with the following parameters: FSH: A, H, W, Voltage 255; SSC: A, H, W, Voltage 203; A488: A, Voltage 198; PE: A, Voltage 341. For each FACS sample 30.000 cells were investigated. As negative control unstained EHD2 del/+ and EHD2 del/del adipocytes were examined at first and the obtained BODIPY intensity values were used as a reference for unstained cells. To exclude adipocytes which did not show any positive fatty acid uptake, all unstained cells were removed resulting in an only positive stained population (R1, illustrated in red in Fig. 2F). Within this R1 population adipocytes with strongly increased BODIPY intensity values were gated to population R2 (blue, Fig. 2F). Detailed analysis/gating and statistics was done by using FlyingSoftware2.5.1 (Perttu Terho, Cell Imaging Core, Turku Center for Biotechnology). For each experiment 15.000 cells were analyzed and gated to the unstained, R1 or R2 population. Next, the percentage of the cells gated to the populations were calculated for every time point and illustrated in the bar graph (Fig. S4C). R2 population was investigated in more detail by normalization to the R2 cell number of EHD2 del/+ adipocytes (Fig. 2H).

#### Glucose uptake assay

Glucose uptake of EHD2 del/+ and EHD2 del/del adipocytes was measured as described by BioVison (2-NBDG Glucose uptake assay). Briefly, adipocytes were treated as described for fatty acid uptake assay. However, after starvation 200 μM 2-NBDG (2-deoxy-2-[(7-nitro-2,1,3-benzoxadiazol-4-yl) amino]-D-glucose, molecular probes #N13195) diluted in serum-free DMEM + 10 μg/ml insulin was applied to the cells followed by incubation times from 5-60 min. Staining analysis was done as mentioned for fatty acid uptake with the same FACS parameters and gating procedure whereby only one positive stained cell population was examined (R1, illustrated in Fig. S4E).

#### Gene expression analysis

EHD2 del/+ and EHD2 del/del adipocytes were differentiated for 5 days, washed twice with ice-cold PBS and RNA was isolated accordingly to the Qiagen protocol (RNeasy Mini Kit, Qiagen). SuperscriptIII First Strand Synthesis Kit (Invitrogen #18080051) was used to obtain corresponding cDNA, which then was used for real-time PCR. Gene expression levels were analyzed by GoTaq q-PCR (Promega, #A6001) Master Mix in Fast real time PCR cycler (Applied Biosystems) accordingly to instructor's protocol. To measure the relative fold change of genes in EHD2 del/del adipocytes compared to EHD2 del/+, the comparative real-time PCR method was applied whereby actin was used as reference gene.

Total mRNA from murine gonadal adipose tissue (gWAT) was extracted with RNeasy Mini Kit (QIAGEN GmbH, Hilden) according to manufacturer's instructions. RNA was transcribed using the Moloney Murine Leukemia Virus Reverse Transcriptase (M-MLV RT, Promega) according to manufacturer's recommendations. Expression of mRNA was determined by quantitative real-time PCR on LightCycler 480 II/384 (Roche, Rotkreuz, Switzerland) using GoTaq Probe qPCR Master Mix (Promega, Madison, USA) applying TaqMan Gene Expression Assays. Target gene expression of was normalized to the mean expression of *Eef2, Ppia* and *Actb* in murine samples.

| Gene | Description | TaqMan Assay |
| --- | --- | --- |
| *EHD2* | EH-domain containing 2 | Hs.PT.58.4969281 |
| *Ehd2* | EH-domain containing 2 | Self-designed |
| *Actb* | Actin, beta | Self-designed |
| *Eef2* | Eukaryotic translation elongation factor 2 | Self-designed |
| *Ppia* | Peptidylprolyl isomerase A | Mm.PT.39a.2.gs |

#### Lipid composition

The measurement of lipid amount and its composition in tissue samples or cells were performed by Lipotype GmbH (Dresden, Germany). For this, tissue samples were homogenized accordingly to the supplied Lipotype protocol and diluted samples (1 mg/ml) were frozen and analyzed by Lipotype. MEF were split and the cell pellet was diluted in cold PBS to a final cell number of 30,000 cells/ml.

#### Statistical analysis

At first, a normality distribution test (Kolmogorov-Smirnov test) was carried out for all experimental values. If the data was normally distributed, Student t-Test (two-tailed P-value) was applied, otherwise Mann-Withney-Rank-Sum (two-tailed P-value) test was used to calculate the significant difference between two groups. Two-way-Anova tests were used to investigate LD size after CD36 siRNA knockdown, whereby for EHD2 wt and KO MEFs each CD36 siRNA#1-3 treated cells were compared to nonsense siRNA (negative control, Fig. 5I). Box plots, if not otherwise indicated in the figure legends, always represents median with whiskers from minimum to maximum, column bar graphs and line graphs represent mean with mean standard error of the mean (SE). Statistical calculations were carried out by using Prism (GraphPad software). Distribution of LD sizes represented in histograms were also obtained by using Prism. For all experiments including the examination of mice or mouse tissue or human samples, n represents the number of mice/patients which were used (Fig. 1, 3, 6, S1–5) and all analyzed cryo/paraffin sections or caveolae are also indicated (e.g.: n = 80 caveolae/3 mice). In cell culture experiments (Fig. 2, 4, 5, S4–7), n represents the number of investigated events (e.g.: lipid droplet area) and the number of independently performed experiments (e.g.: n = 80 lipid droplets/3 independent experiments). The following P-values were used to indicate significant difference between two groups: * P<0.05; ** P<0.001; *** P<0.0001.

## Supplemental Material

**Movie S1: Electron tomography of EHD2 del/del BAT (related to Fig. 3)**

Caveolae in a 150 nm EHD2 del/del BAT section were investigated by electron tomography. The movie includes all images obtained during the tilting from – 60° to 60° (image acquisition every 2°). Both, detached and plasma membrane bound caveolae can be observed. For better handling during the segmentation and reconstruction in IMOD, the tomogram was reflected horizontal.

**Movie S2-3: TIRF live imaging of caveolae in EHD2 +/+ and del/del MEFs (related to Fig. 4)**

EHD2 +/+ and del/del MEFs were transfected with pCav1-EGFP to detect single caveolae and afterwards TIRF live imaging was performed. Movie S2 illustrates an example EHD2 +/+ MEF, and Movie S3 shows an EHD2 del/del MEF. Notably, Cav1 spots in MEFs lacking EHD2 showed increased caveolar dynamics.

**Movie S4: TIRF live imaging of caveolae in EHD2 del/del MEFs transfected with pEHD2-EGFP (related to Fig. 4)**

EHD2 del/del MEFs were co-transfected with pCav1-EGFP and pEHD2-EGFP and caveolae movement was observed by TIRF live imaging. Re-expression of EHD2 in EHD2 del/del MEFs strongly reduced the dynamics of caveolae.

**Fig. S1:**
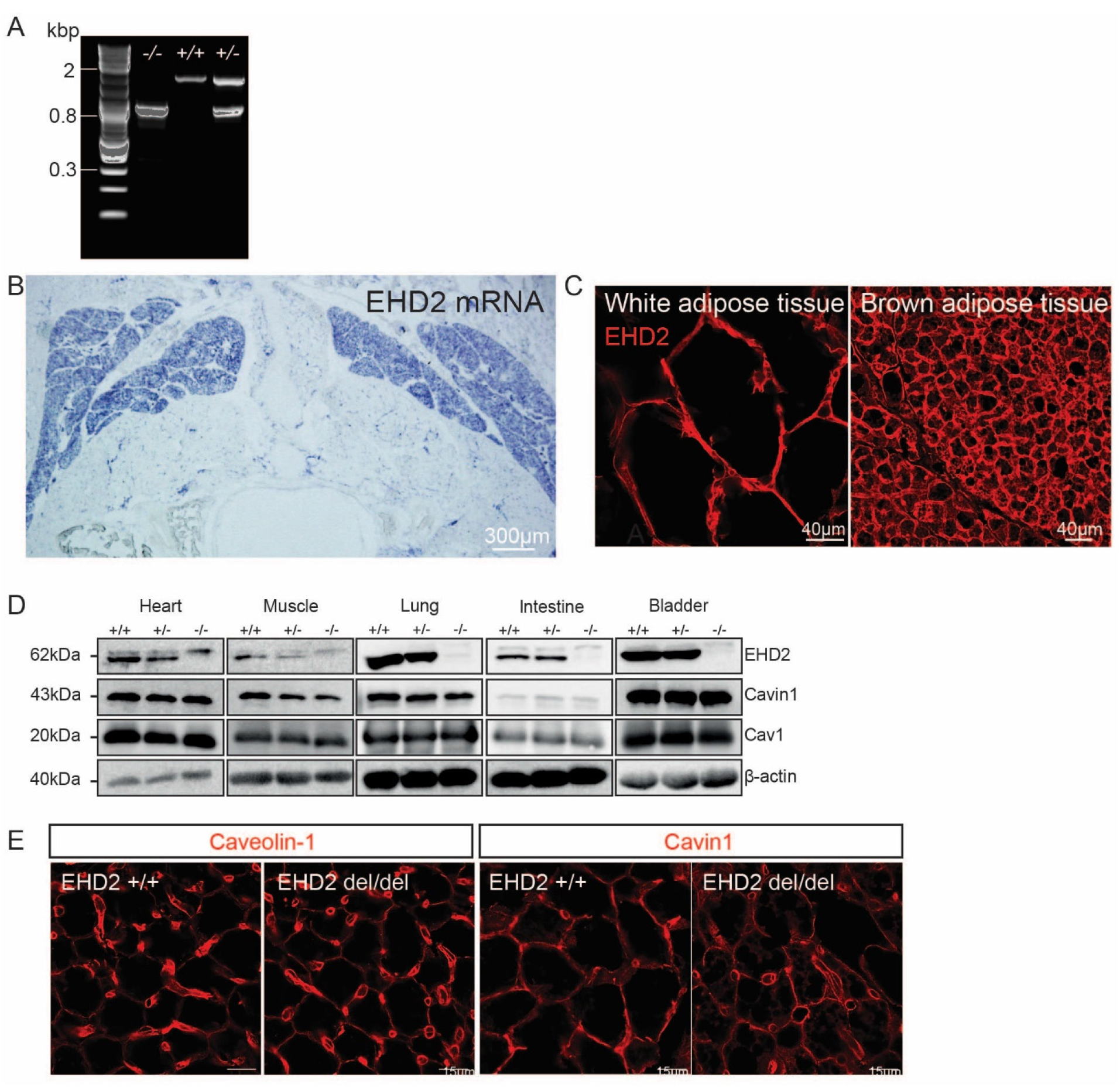
EHD2, Cav1 and Cavin1 expression in BAT (related to Fig. 1) **A** Genotyping of EHD2 delta E3 offspring (wildtype – band size 1,700 bp; EHD2 KO – band size 830 bp). **B** *In situ* hybridization against EHD2 mRNA of BAT in an E18 C57BL6/N embryo. **C** Cryostat section of adult C57BL6/N white adipose or brown adipose tissue stained against EHD2. **D** Western Blot analysis of different tissues from EHD2+/+, +/− and −/− mice showing EHD2, Cav1 and Cavin1 protein level. **E** Cavin1 and Cav1 protein level in BAT cryostat sections from EHD2 wt and del/del mice.

**Fig. S2:**
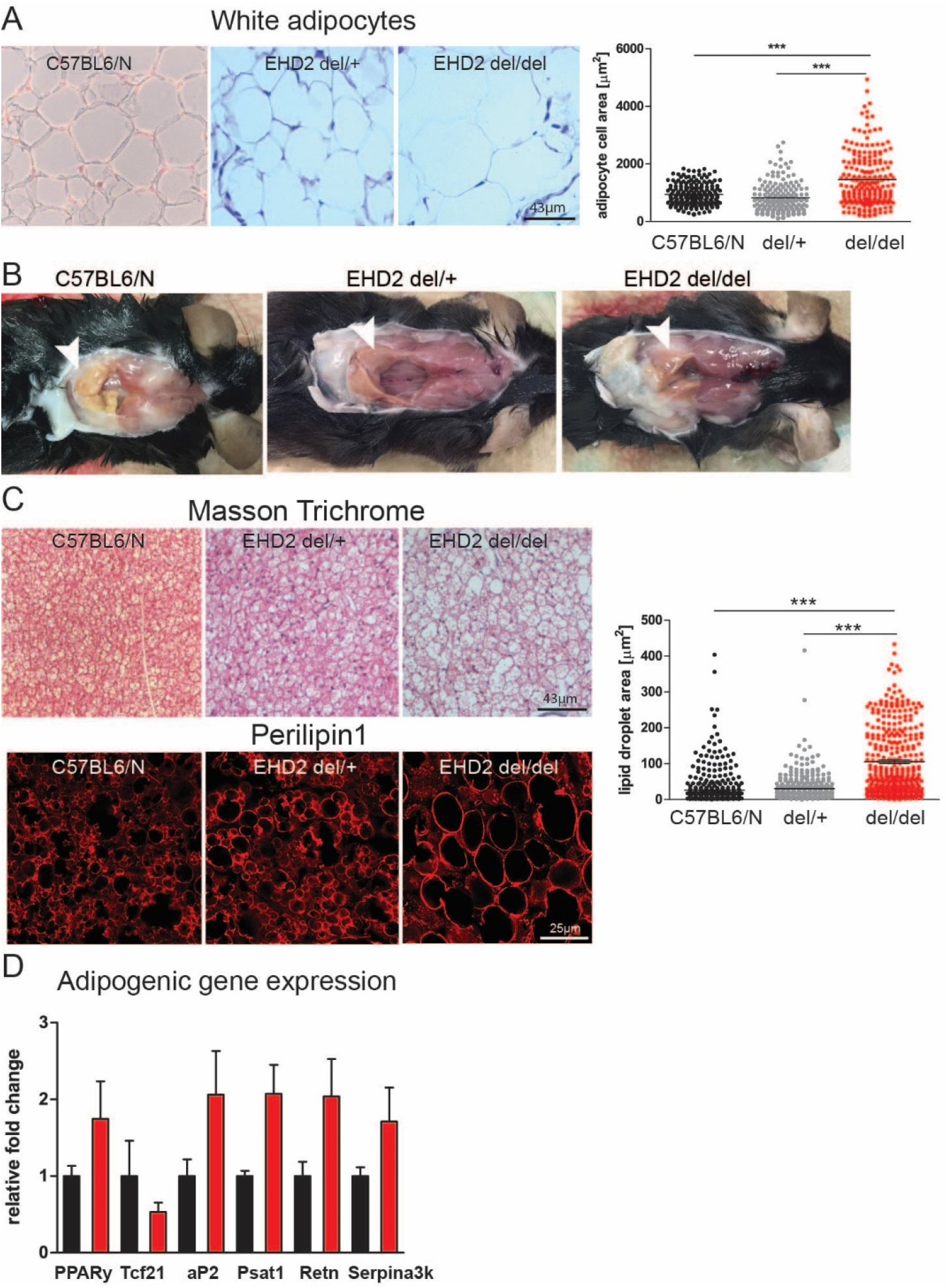
C57BL6/N and EHD2 del/+ mice did not reveal any differences in lipid accumulation (related to Fig. 1) **A** WAT paraffin sections from C57BL6/N mice stained with Masson Trichrome and analyzed by adipocyte cell size (n(C57BL6/N) = 186/3; n(EHD2 del/+ = 172/3; n(EHD2 del/del) = 199/3). **B** BAT examples of C57BL6/N, EHD2 del/+ and del/del mice. **C** BAT paraffin and cryostat sections stained against Perilipin1 obtained from C57BL6/N mice (lipid droplet size was measured by Perilipin1 staining, n(C57BL6/N) = 461/3; n(EHD2 del/+ = 398/3; n(EHD2 del/del) = 352/3). **D** Adipogenic gene expression analysis of WAT from EHD2 del/+ and EHD2 del/del mice (n = 8). Graphs illustrate each replicate with mean +/− SE, column bar graphs show mean + SE, normal distributed groups were analyzed by t-test, not normally distributed values with Mann Withney U test, * P<0.05, *** P<0.0001.

**Fig. S3:**
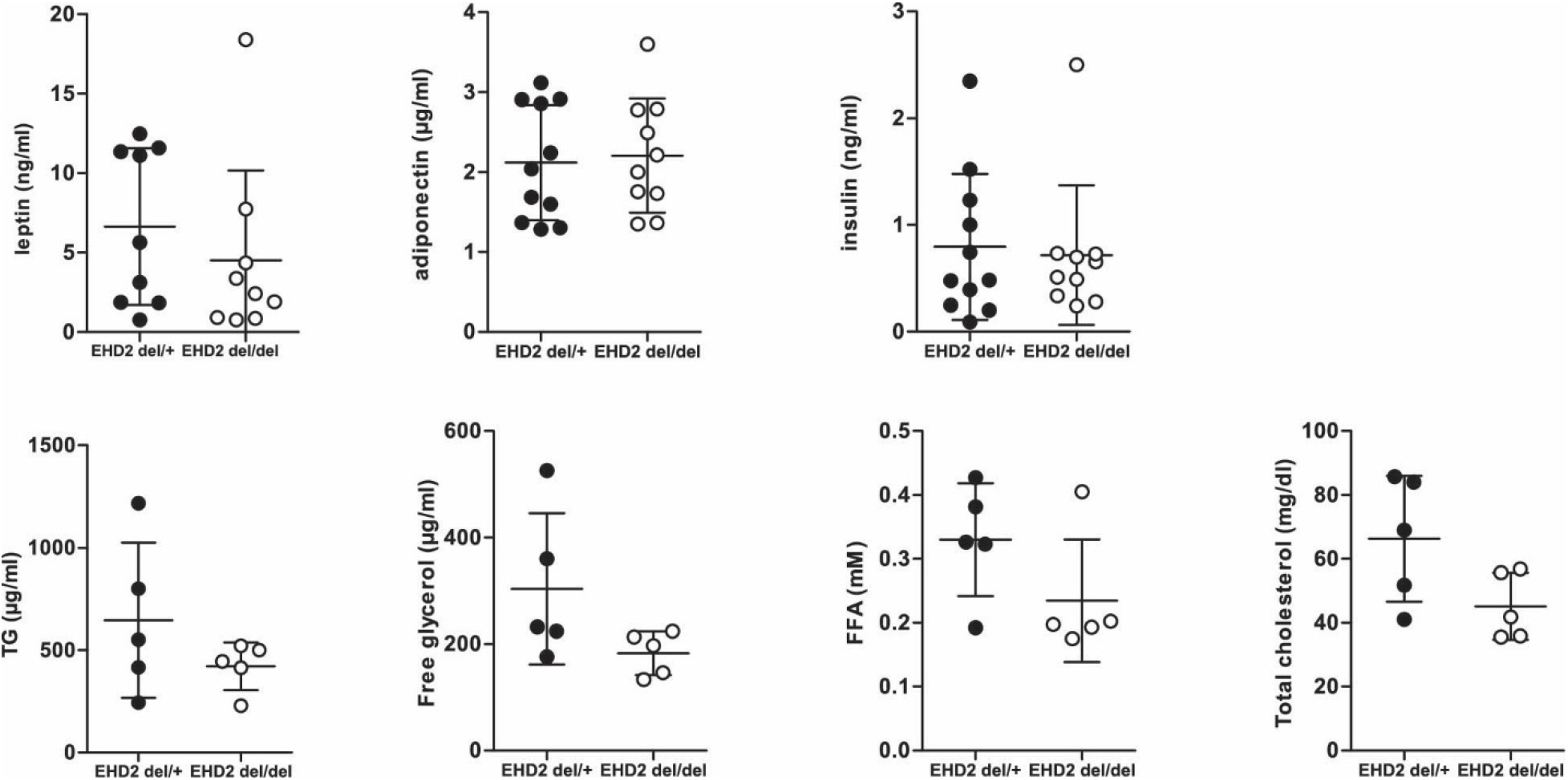
Blood plasma analysis of EHD2 del/+ and EHD2 del/del mice did not reveal any significant differences (related to Fig. 1) Blood samples obtained from EHD2 del/+ and del/del mice (n(EHD2 del/+) = 10 or 5; n(EHD2 del/del) = 10 or 5; graph illustrates each replicate with mean +/− SE). FFA – free fatty acid, TG – triglycerol.

**Fig. S4:**
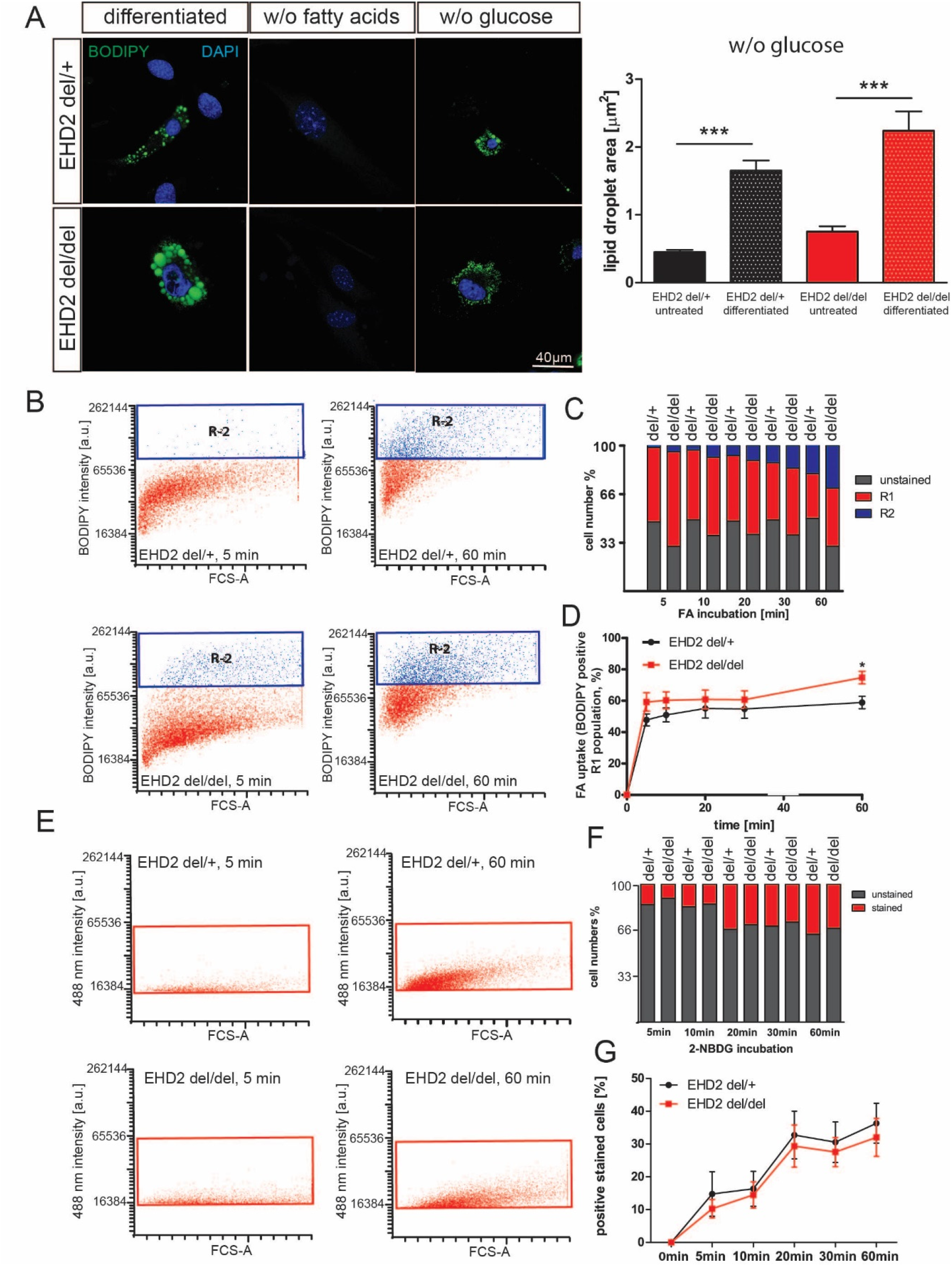
Cellular investigation of EHD2 del/del adipocytes (related to Fig. 2) **A** Pre-adipocytes were treated with either differentiation medium containing delipidated FBS or without glucose and BODIPY staining illustrating LDs. **B-D** Fatty acid uptake assay in differentiated EHD2 del/+ and EHD2 del/del adipocytes. Dodecanoic acid-BODIPY uptake was measured after 5, 10, 20, 30 or 60 min, and R1 population indicates positively stained cells (illustrated in red in graphs B, C). R2 populations (blue) correspond to higher BODIPY staining intensity in cells and represent adipocytes with increased amount of dodecanoic acid taken up (shown in blue in graphs B, C). Overview of fatty acid uptake (percent cell numbers (B), and time scale (D); B-D, n(del/+) = 6/3 experiments, n(del/del) = 8/3 experiments). **E-G** Following 5 days of differentiation, glucose uptake in cultured adipocytes was measured after 5-60 min (C, n = 6). Line graphs show mean +/− SE, column bar graphs show mean + SE, t-test or Mann Withney U test was used to calculate significance, ** P<0.001, *** P<0.0001.

**Fig. S5:**
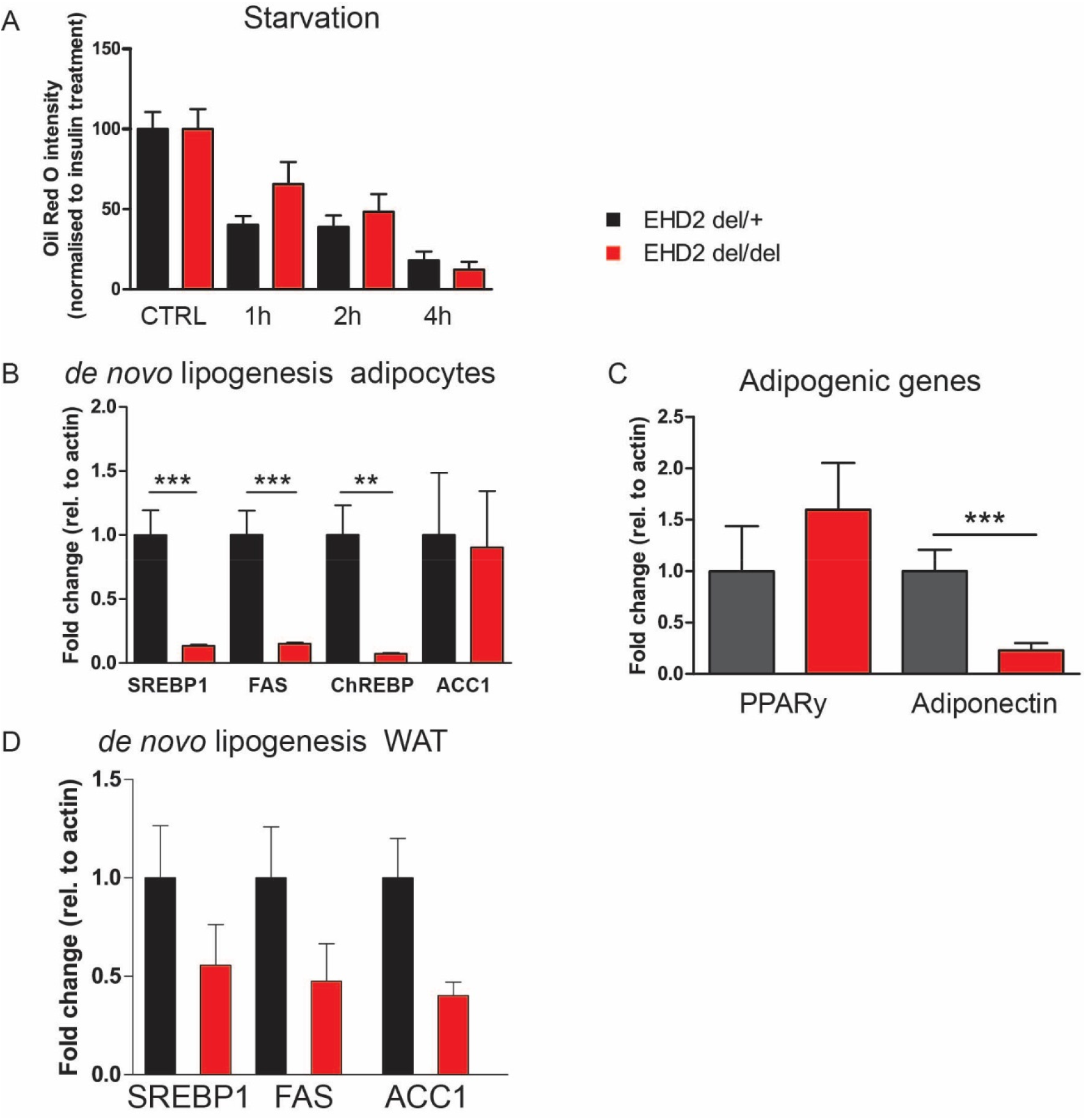
Cellular investigation of EHD2 del/del adipocytes (related to Fig. 2) **A** Differentiated EHD2 del/+ and EHD2 del/del adipocytes were starved for 1-4h and Oil Red O staining was applied (n(EHD2 del/+ = 53/3; n(EHD2 del/del) = 50/3). **B-C** Gene expression analysis of genes involved in *de novo* lipogenesis (B) or adipogenic genes (C) in EHD2 del/+ and del/del differentiated adipocytes (n = 8). **D** Gene expression analysis of genes involved in *de novo* lipogenesis (n = 5). Column bars show mean + SE, t-test or Mann Withney U test was used to calculate significance, ** P<0.001, *** P<0.0001.

**Fig. S6:**
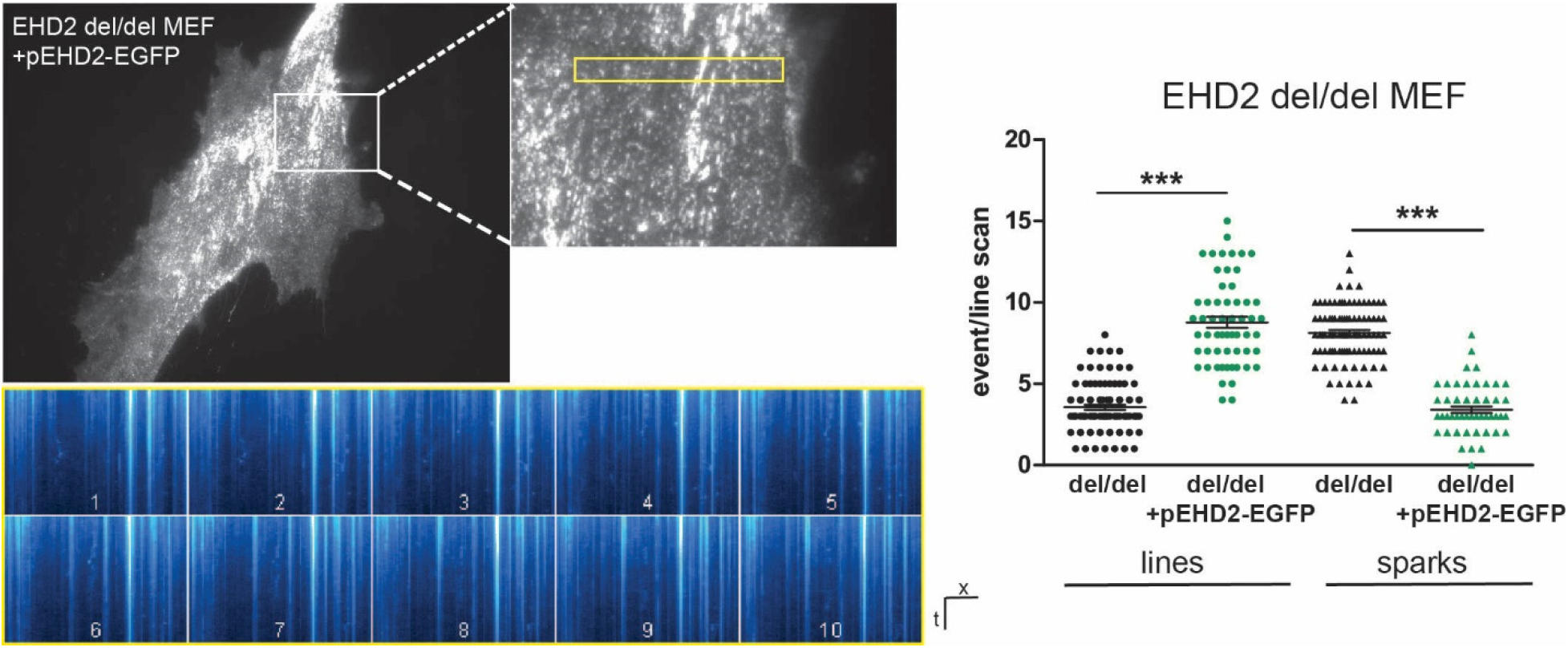
Caveolar mobility in EHD2 del/del MEFs (related to Fig. 4) TIRF live-imaging in EHD2del/del MEFs transfected with pEHD2-EGFP and pCav1-EGFP (1:1) to investigate single caveolae movement by TIRF live imaging. Non-moving caveolae correspond to vertical lines within the line scan, moving caveolae can be related to single sparks (B, n = 30, graph illustrates each replicate with mean +/− SE, t-test was used to calculate significance). See also Movie S4. Plots indicate each replicate from maximal to minimum value with mean, t-test or Mann Withney U test were used to calculate significance, *** P<0.0001.

**Fig. S7:**
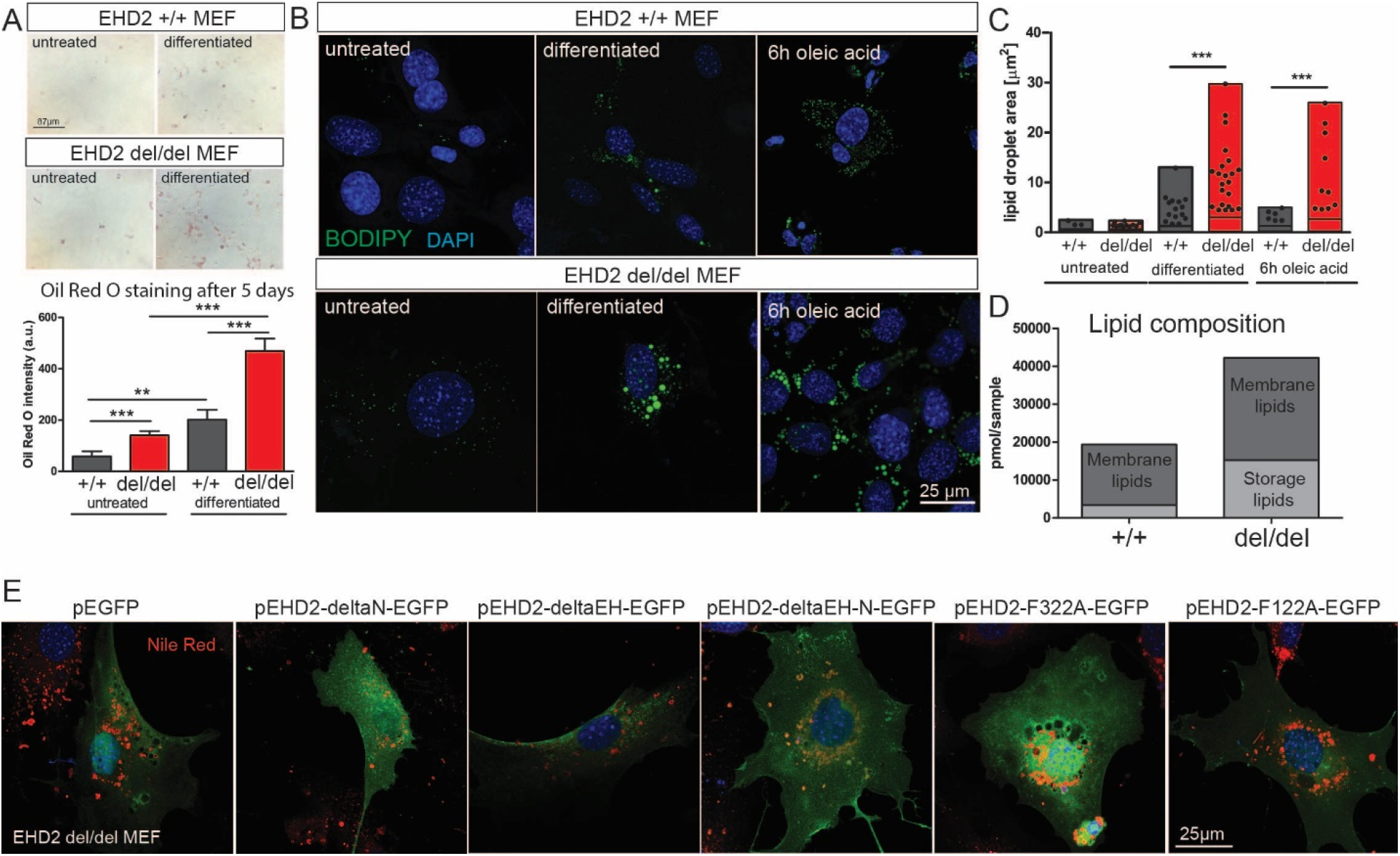
Caveolar mobility in EHD2 del/del MEFs (related to Fig. 5) **A** Oil Red O staining of EHD2 +/+ and del/del MEFs untreated or treated with adipocyte differentiation medium (untreated: n(+/+) = 21/3, n(del/del) = 28/4); differentiated: n(+/+) = 37/3, n(del/del) = 55/4). **B-C** LD analysis after 5 days of differentiation or 6 h of oleic acid in EHD2 MEFs by BODIPY staining (D, untreated: n(+/+) = 61/2, n(del/del) = 139/4); differentiated: n(+/+) = 148/3, n(del/del) = 200/4; oleic acid: n(+/+) = 146/3, n(del/del) = 217/3). **D** Lipid composition analysis of MEFs differentiated for 5 days, 120,000 cells were analyzed for each experiment (n = 3). **E** EHD2 del/del MEFs were transfected with different EHD2 constructs and lipid droplet size was analyzed after 6h oleic acid treatment. Box plots indicate each replicate from maximal to minimum value with mean, column bar graphs show mean + SE, t-test or Mann Withney U test were used to calculate significance, ** P<0.001, *** P<0.0001.

